# Explainable Machine Learning Identifies Factors for Dosage Compensation in Aneuploid Human Cancer Cells

**DOI:** 10.1101/2025.05.12.653427

**Authors:** Erik Marcel Heller, Karen Barthel, Markus Räschle, Klaske M. Schukken, Jason M. Sheltzer, Zuzana Storchová

## Abstract

Aneuploidy, a hallmark of cancer, leads to widespread changes in chromosome copy number, altering the abundance of hundreds or thousands of proteins. However, evidence suggests that levels of proteins encoded on affected chromosomes are often buffered toward their abundances observed in diploid cells. Despite its prevalence, the molecular mechanisms driving this protein dosage compensation remain largely unknown. It is unclear whether all proteins are buffered to a similar degree, what factors determine buffering, and whether dosage compensation varies across different cell lines or tumor types. Moreover, its potential adaptive advantage and therapeutic relevance remain unexplored. Here, we established a novel approach to quantify protein dosage buffering in a gene copy number-dependent manner, showing that dosage compensation is widespread but variable in cancer cell lines and *in vivo* tumor samples. By developing multifactorial machine learning models, we identify mean gene dependency, protein complex participation, haploinsufficiency, and mRNA decay as key predictors of buffering. We also show that dosage compensation can affect oncogenic potential and that higher buffering correlates with reduced proteotoxic stress and increased drug resistance. These findings highlight protein dosage compensation as a crucial regulatory mechanism and a potential therapeutic target in aneuploid cancers.

## Introduction

Chromosome and chromosome arm copy number changes, commonly referred to as chromosomal aneuploidy, are a widespread hallmark of cancer (Santaguida & Amon, 2015; Sdeor et al., 2024; Taylor et al., 2018). Aneuploidy alters the expression of many genes located on the affected chromosomes, including tumor suppressors and oncogenes. The altered gene expression disrupts cellular regulation, driving tumorigenesis and promoting cancer progression. This likely explains why certain chromosomes or chromosome arms are recurrently gained or lost across various cancers and why specific aneuploidies are frequently associated with distinct cancer types (Davoli et al., 2013; Shih et al., 2023). For example, trisomy 12 is common in chronic lymphocytic leukemia, while loss of chromosome 13q is prevalent in retinoblastoma. Aneuploidy is often associated with chromosomal instability and together they facilitate tumor evolution by providing selective advantage to cancer cells, increasing drug resistance, and metastatic dissemination (Andrade et al., 2023; Lukow & Sheltzer, 2022; Replogle et al., 2020; Shukla et al., 2020).

However, aneuploidy affects the copy number of hundreds of genes, and this deregulated gene expression may have negative effects on cellular proliferation. Evidence from *in vitro* models of aneuploidy in budding yeast, murine and human cells revealed that aneuploidy often impairs proliferation, induces cellular senescence, increases genomic instability, and disrupts metabolic homeostasis. Remarkably, aneuploid cells exhibit increased sensitivity to compounds that interfere with protein folding and turnover (Donnelly & Storchová, 2015; Donnelly et al., 2014; Tang et al., 2011; Torres et al., 2010). This suggests that the aberrant gene expression in aneuploid cells overwhelms cellular quality control systems involved in the maintenance of protein homeostasis. Indeed, aneuploid cells often experience an imbalance in the synthesis and degradation of proteins, leading to the accumulation of misfolded proteins, which must be resolved by the proteasome and other protein degradation pathways (Donnelly & Storchová, 2015; Donnelly et al., 2014; Oromendia & Amon, 2014; Oromendia et al., 2012). Moreover, secondary global (i.e., *in trans*) gene expression changes are often observed in aneuploid cells, as gene expression of multiple genes across the entire genome is altered in response to the stresses induced by aneuploidy (Sheltzer et al., 2012; Stingele et al., 2012).

Transcriptome and proteome analysis of engineered aneuploid cells have provided useful insights into the gene expression changes triggered by aneuploidy. For instance, in budding yeast strains with an extra chromosome, most genes on the aneuploid chromosome show an increased abundance of both mRNA and proteins, reflecting the proportional increase in gene dosage (Dephoure et al., 2014; Torres et al., 2007), but the abundance of approximately ∼20% of proteins encoded on extra chromosomes shows dosage compensation, meaning the protein levels are adjusted to levels similar to euploid levels (Figure 1A; Dephoure et al., 2014). This dosage compensation, also referred to as protein abundance attenuation or buffering, is particularly apparent for subunits of multi-protein complexes and largely depends on proteasomal and lysosomal degradation of the excess proteins, and their sequestration into aggregates (Brennan et al., 2019; Dephoure et al., 2014; Senger & Schaefer, 2021; Taggart et al., 2020). In naturally occurring aneuploid yeast strains, the dosage compensation affects an even larger fraction of proteins and is accompanied by enhanced protein turnover via proteasomal degradation (Muenzner et al., 2024). Analysis of engineered cancerous and non-cancerous human cell lines harboring chromosome gains demonstrated that most genes on gained chromosomes are transcribed and translated proportionally to the chromosome copy number gain. However, protein complex subunits, protein kinases, and several other proteins showed an apparent dosage compensation at the protein level (Stingele et al., 2012; Viganó et al., 2018). Similarly, dosage compensation modulates the abundance of proteins in cells from individuals with trisomy syndromes (Hwang et al., 2021; Liu et al., 2017). Loss of a single chromosome copy is also compensated in engineered monosomic human cells (Chunduri et al., 2021). How dosage changes of chromosomes and single genes shape the proteome in human cancers is poorly understood. Gene amplification often results in increased protein abundance, but in up to ∼30% of cases the abundance changes do not scale linearly (Gonçalves et al., 2017; Mertins et al., 2016; Sousa et al., 2019; B. Zhang et al., 2014; H. Zhang et al., 2016). Recently, the Cancer Cell Line Encyclopedia (DepMap CCLE) provided data for hundreds of cell lines with matched DNA copy number, RNA expression, and protein expression measurement levels (DepMap, Broad Institute, 2023; Ghandi et al., 2019; Nusinow et al., 2020). Analysis of this dataset revealed that the mRNA abundances of the majority of genes scale with gene dosage changes, while protein abundance frequently changes less than expected based on the chromosome copy number (Schukken & Sheltzer, 2022). Thus, aneuploidy is subject to dosage compensation at the protein level even in cancer cell lines. The existence of dosage compensation has been demonstrated in aneuploid ovarian tumors (Schukken & Sheltzer, 2022). However, a comprehensive overview of dosage compensation across multiple *in vivo* tumor types remains to be established. Previous studies by Schukken & Sheltzer aimed to identify the mechanisms involved in dosage compensation, but the predictive quality of individual factors analyzed in the study was rather low (Schukken & Sheltzer, 2022).

**Figure 1:**
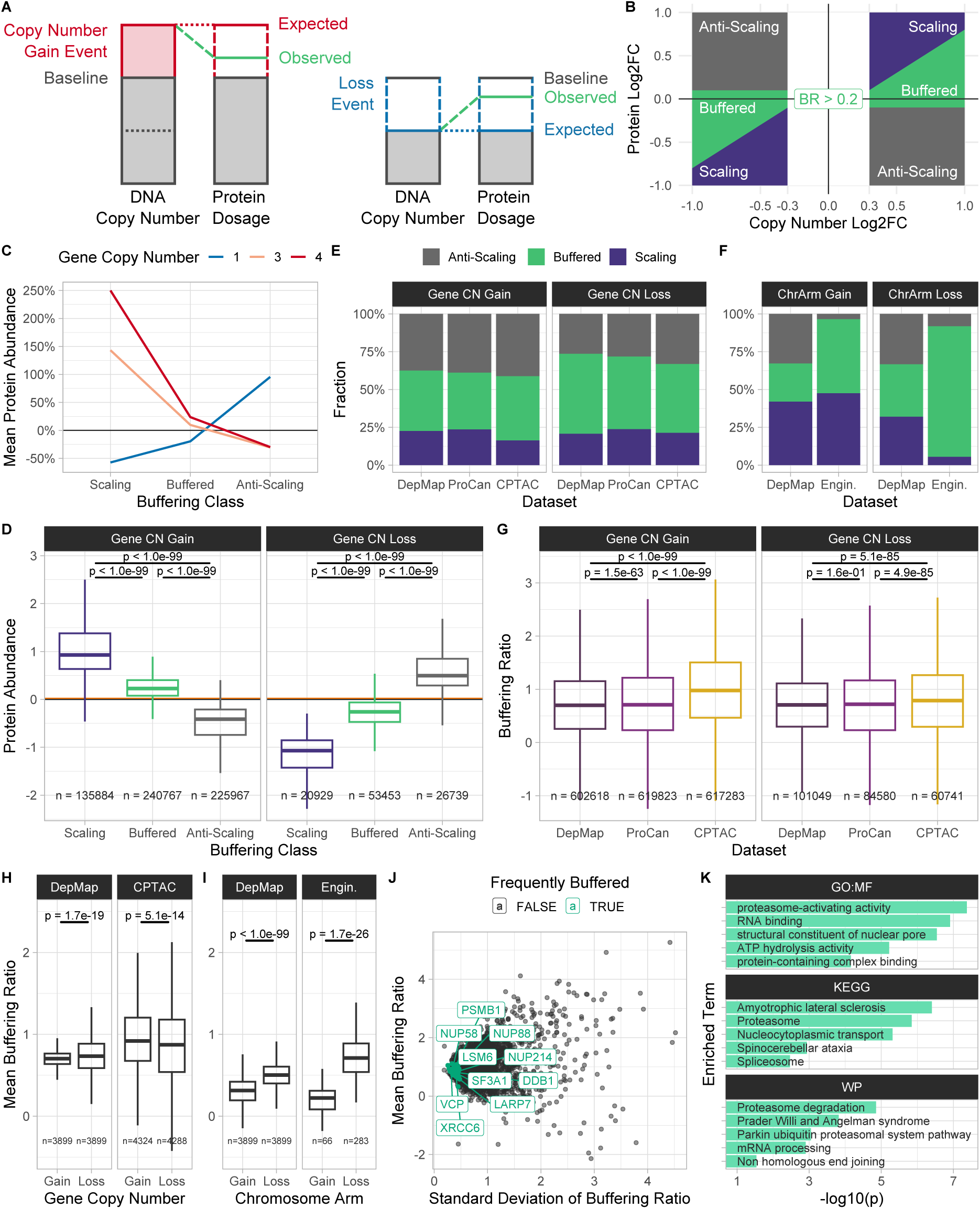
Quantification of protein dosage compensation in aneuploid cancer cells. **(A)** Illustration of protein dosage compensation upon gene copy number gain (left) and loss (right). **(B)** Illustration of the buffering classes “Scaling”, “Buffered”, and “Anti-Scaling”. A threshold on the buffering ratio (BR) was used to distinguish “Buffered” (green) from “Scaling” (indigo) observations, while protein log2 fold-change thresholds were used to distinguish “Buffered” and “Anti-Scaling” (grey) observations. **(C)** Mean change in protein abundance relative to protein abundance baseline for proteins in each buffering class in the DepMap CCLE pan-cancer cell line dataset. **(D)** Difference in pro-tein abundance between the different buffering classes (shown dataset: DepMap). The mean disomic protein abundance baseline is represented as an orange line. **(E)** Categorical distribution of buffering classes across pan-cancer datasets (DepMap, ProCan, CPTAC) upon gene copy number gain and loss. **(F)** Categorical distribution of buffering classes across DepMap CCLE and lab-engineered cell lines upon chromosome arm gain and loss. **(G)** Difference in buffering ratio distribution of all proteins with either gene copy number gain or loss between pan-cancer datasets (DepMap, ProCan, CPTAC). **(H)** Difference in buffering ratio distribution upon gene copy number gain and loss across pan-cancer datasets (DepMap, CPTAC). **(I)** Difference in buffering ratio distribution upon chromo-some arm gain and loss across DepMap CCLE and lab-engineered cell lines. **(J)** Mean and standard deviation of buffering ratio of proteins across samples from the pan-cancer tumor sample dataset CPTAC. Highlighted are 50 genes with lowest buffering ratio vari-ance among frequently buffered genes across all pan-cancer datasets (DepMap, ProCan, CPTAC). **(K)** Top 5 enriched terms per selected data source in over-representation ana-lysis (ORA) of highlighted low-variance genes.

Here, we present a comprehensive analysis of protein dosage compensation in human samples by developing a novel approach that quantifies compensation at single-gene resolution across copy number changes of somatic chromosomes. Exploiting pan-cancer cell line and tumor sample datasets from DepMap CCLE, ProCan, CPTAC, and lab-engineered aneuploid cells lines, our method quantifies dosage compensation for each gene within single samples using focal gene and chromosome-wide copy number alterations, providing increased granularity. By integrating diverse biochemical and genetic variables from multiple resources into explainable machine learning models, we achieved high predictive accuracy, uncovering critical mRNA- and protein-level features that drive compensation of aneuploid somatic chromosomes. Notably, our findings reveal that cancers with low dosage compensation exhibit more vulnerabilities than those with high compensation — a disparity that can be exploited for targeted therapies. Overall, our results position dosage compensation as a factor contributing to tumorigenesis and cancer resilience.

## Results

### Calculating protein-level attenuation using absolute gene copy numbers in human cell lines and tumor samples

Precise quantification of dosage compensation relies on the analysis of copy number changes across the entire genome matched to the changes observed on the proteome level. In previous studies, copy number changes were commonly evaluated as gains or losses of entire chromosomes or chromosome arms. However, we frequently observed sub-arm copy number changes in tumor tissues and cancer cells (Figure S1A). To increase the granularity of such analyses, we derived a novel indicator we term Buffering Ratio (BR) which compares both focal gene and chromosome-wide copy number changes against changes in protein abundance. The BR is defined as the difference between protein abundance log2 fold-change (Log2FC) and (gene or chromosome) copy number Log2FC (see Methods). This definition allows the use of numeric gene and chromosome arm copy number differences for buffering quantification, thus increasing the precision of the buffering estimate, compared to *a priori* categorization of chromosome arm copy number alterations (CNAs) into gain, neutral, and loss as previously applied (Schukken & Sheltzer, 2022). The BR also allows protein buffering quantification for each gene within each sample (cell line or tumor sample), thus improving analysis granularity compared to methods that average protein buffering per gene across a set of samples.

Based on the BR and the co-directionality of the protein abundance changes with the gene copy number, we categorized genes into the buffering classes *Scaling*, if the change in protein abundance was proportional to the change in gene copy number, *Buffered* if the protein abundance change was less than proportional, and *Anti-Scaling* if these values were anti-proportional (Figure 1B). *Scaling* and *Buffered* proteins were distinguished using a threshold on the copy number-dependent BR, whereas *Buffered* and *Anti-Scaling* proteins were differentiated using a copy number-independent protein abundance Log2FC threshold. We calculated the BR and buffering classes for each gene and sample using chromosome arm CNAs (Cohen-Sharir et al., 2021) and absolute gene copy numbers (Carter et al., 2012; DepMap, Broad Institute, 2023) for the pan-cancer cell line datasets DepMap CCLE containing proteomics data for 375 cell lines, Sanger ProCan with 949 cell lines, and to the pan-cancer tumor sample dataset CPTAC with 1026 tumor samples and 523 normal tissue samples. We removed tumor samples with a purity below 40% to control for tumor purity. Additionally, we analyzed near-diploid p53-deficient human retinal pigment epithelium cell lines (RPE-1) engineered to carry either monosomies or trisomies on single chromosomes (Chunduri et al., 2021; and data from this study). As expected, proteins classified as *Scaling* using the BR showed a change in protein abundance proportional to the change in copy number, whereas *Buffered* proteins retained near-disomic levels (Figure 1C&D). Across datasets, we found that the majority of proteins were either *Buffered* (38%-53%) or *Anti-Scaling* (26%-41%) upon gene copy number gain and loss across pan-cancer cell line (DepMap, ProCan) and tumor sample datasets (CPTAC; Figure 1E). CPTAC had the highest fraction of *Buffered* proteins upon gene copy number gain (42%), and DepMap had the highest fraction of *Buffered* proteins upon gene copy number loss (53%). Additionally, the fractions of *Buffered* observations were higher upon gene copy number loss than upon gain (gain: 38%-42%, loss: 45%-53%), whereas *Anti-Scaling* was reduced upon copy number loss (gain: 37%-41%, loss: 26%-33%). This pattern was preserved in the pan-cancer cell line dataset DepMap CCLE when using chromosome arm CNA data (Figure 1F), and was also preserved when controlling for whole genome doubling (WGD; Figure S1C). Interestingly, lab-engineered cell lines showed less *Anti-Scaling* proteins compared to pan-cancer datasets and almost all proteins affected by chromosome arm loss were classified as *Buffered* (Figure 1F, Supplementary Table 1). Thus, gene dosage buffering is a widespread phenomenon in engineered aneuploid cells.

**Table 1:** Factors used for the dosage compensation factor analysis grouped by data source. Preprocessing describes how datasets have been transformed to be used as factors.

| Factors | Preprocessing | Source |
| --- | --- | --- |
| Protein-Protein Interactions | Counted number of interactions per protein | HIPPIE, Version: v2.3<br>(Alanis-Lobato et al., 2017) |
| Protein Complexes | Counted number of complexes a protein participates in | CORUM<br>Version: 210512<br>(Giurgiu et al., 2019) |
| Mean 3'-UTR Length,<br>Mean 5'-UTR Length | Difference between transcription start<br>and coding region start (5'-UTR), and<br>coding region end and transcription<br>end (3'-UTR), averaged per gene | UCSC Human Gen-<br>ome Browser<br>Track: NCBI RefSeq<br>Assembly:<br>GRCh38/hg38<br>(UCSC Genome<br>Bioinformatics Group,<br>n.d.) |
| Phosphorylation, Ubi-<br>quitination, Sumoyla-<br>tion, Methylation,<br>Acetylation, and Reg-<br>ulatory Sites; Kinase<br>Interactions | Counted number of sites / interactions<br>per protein | PhosphoSitePlus<sup>®</sup>,<br>www.phosphosite.org<br>(Hornbeck et al.,<br>2015) |
| mRNA/Protein<br>Abundance, Tran-<br>scription/Translation<br>Rate, Protein/mRNA<br>Length | – | humanRates.ods<br>(Hausser, 2019) |
| Intrinsic Protein Dis-<br>order, Low Complex-<br>ity Score, Homology<br>Score, Loops In Pro-<br>tein Score, Protein<br>Polyampholyte Score,<br>Protein Polarity | – | MobiDB<br>Version: 5.0<br>Release: 2022_07<br>Proteome:<br>UP000005640<br>(Piovesan et al., 2021) |
| Non-Exponential De-<br>cay Delta | Mean per protein | Supp. Table 4<br>(McShane et al., 2016) |
| Mean mRNA Decay<br>Rate | Mean per gene | Supp. Table 9 (Yang<br>et al., 2003) |
| Protein Half-Life | Mean per gene | Supp. Table 2 (Math-<br>ieson et al., 2018) |
| Aggregation Score | – | Supp. Table 2<br>(Ciryam et al., 2013) |
| Haploinsufficiency /<br>Triplosensitivity Score | – | Table S7 (Collins et<br>al., 2022) |
| Mean Gene Depend-<br>ency | Mean dependency score per gene | DepMap (DepMap,<br>Broad Institute, 2023) |
| Transcription Factors (Repressor/Activator/All), Mean TF Regulation Mode | Counted the number of transcription factors (TFs) per gene. TFs with a negative mode of regulation (MOR) were counted as repressors, TFs with positive MOR as activators. Calculated the mean MOR per gene. | Obtained DoRothEA regulons using the **decoupleR** package (levels A,B,C,D). (Badia-i-Mompel et al., 2022; Garcia-Alonso et al., 2019) |
| Random Allelic Expression | Combined male and female z-score columns of hc-RAE and hc-Biallelic (all tissues) and calculated mean per gene | Table S2 (Kravitz et al., 2023) |

Comparing BR distributions between datasets showed that tumor samples have a significantly increased BR upon gain and loss compared to cell lines, in line with the observed increase in *Buffered* and *Anti-Scaling* proteins (Figure 1G). We did not observe a significant correlation between tumor purity and BR that could confound these results (Figure S1D). Instead, we observed slightly increased immune and stromal cell invasion scores in tumor samples with high average BRs (Figure S1D). To understand the changes in buffering in different conditions, we compared the BR distributions between (gene copy number and chromosome arm) gains and losses. We observed that the BR was significantly higher upon loss than upon gain events in lab-engineered cell lines and pan-cancer cell lines, while in tumor samples the BR was higher upon gene copy number gain (Figure 1H&I). This might be caused by the fact that tumor samples harbored increased fractions of genes with strong copy number amplifications (Figure S1A). Therefore, tumor samples may also show increased buffering to restore the protein abundance of these genes to near-disomic levels. Interestingly, the BR was lower upon gene copy number loss than upon gain in cell lines that underwent WGD, but was higher upon loss in Non-WGD cell lines (Figure S1). This suggests that buffering might be especially important if only one gene copy is available. We conclude that more protein buffering occurs upon copy number loss than upon gain events in aneuploid cell lines independent of whether these are chromosomal aberrations or focal gene copy number variations.

Having calculated the buffering ratios per gene and per sample, we asked which genes are frequently and consistently buffered independent of cell states. To do this, we filtered for genes which were classified as *Buffered* in more than 33% of samples in all pan-cancer datasets (DepMap, ProCan, CPTAC) and whose BR had a standard deviation smaller than 2. We then aggregated their BR standard deviation ranks across datasets. In the top 10 frequently buffered genes with low BR variation were NUP88, NUP214, and NUP58, three subunits of the nuclear pore complex; ribonucleoproteins LARP7, SF3A1, and LSM6; DNA repair proteins XRCC6, and DDB1; proteasomal subunit PSMB1; and VCP, a protein that segregates protein molecules from large protein assemblies to facilitate their degradation in the proteasome (Figure 1J). All these proteins are subunits of macro-molecular complexes, which were previously suggested to be subject to extensive dosage compensation (Stingele et al., 2012). The only exception is the segregase VCP/p97, which interacts with many proteins but is not a stable member of a macromolecular complex. Importantly, VCP has a key role in protein quality control, which is often challenged in aneuploid cells (Gwon et al., 2021; Kadowaki et al., 2015; Oromendia & Amon, 2014; Oromendia et al., 2012; Turakhiya et al., 2018). Over-representation analysis (ORA) further confirmed that RNA binding and processing, spliceosome, and proteasome genes were significantly enriched among the top 50 frequently buffered genes (Figure 1K). This confirms the previous notion that subunits of macromolecular complexes need to be maintained at stoichiometric levels.

These results were reproducible, as we observed a 50% mean correlation of gene copy number-derived BRs between the cell line datasets DepMap and ProCan and a 55% mean correlation for the top 50 frequently buffered genes (Spearman; Figure S1E&F).

### Average buffering per sample relates to cellular and clinical fea-tures of cancer

Next, we asked whether protein dosage compensation differs among different cell lines and tumor types. Using the same datasets as previously, we calculated sample buffering ratios (sample BRs) by calculating the mean BR per sample (cell line or tumor sample). Calculating the z-scores of sample BRs showed significant differences in average buffering between cell lines (Figure 2A). These results were consistent across cell line datasets, as the calculated sample BRs ratios were highly correlated between DepMap and ProCan. The high correlation of the BR allowed us to aggregate the sample BR ranks of cell lines across the datasets using the mean normalized rank (MNR, Upadhya & Ryan, 2022) to provide a resource of dataset-independent average buffering estimates for each cell line (Figure 2B, Supplementary Table 3). Plotting the mean sample BR per cancer type and dataset using the OncoTree classification (Kundra et al., 2021) showed differences in average buffering between cancer types (Figure 2C). Among cancer types with low average buffering we observed fibrosarcomas, hepatoblastomas, Erwing sarcomas, neuroblastomas, and myeloid/lymphoid neoplasms, whereas pancreatic neuroendocrine tumors, pleural mesotheliomas, hepatocellular carcinomas, anaplastic thyroid cancers, and invasisve breast carcinomas were among the high-buffering cancer types. In particular, pediatric cancers possessed lower sample BRs on average than adult cancers and that high-buffering tumor samples showed slightly lower patient survival probabilities (Figure S2C&D). Interest-ingly, TP53 mutations had a non-uniform effect, but we observed that the sample BR is positively correlated with the aneuploidy score (AS) and average ploidy of cell lines (Figure 2C&D, Figure S2A,B&E). Furthermore, separating cell lines into cohorts with or without whole genome doubling (WGD) showed significant differences in sample BR between these cohorts suggesting that WGD-positive cells exhibit more protein buffering on average (Wilcoxon rank-sum test, ProCan: *p* = 8.0 ∗ 10*^−^*^44^). We conclude that WGD-positive cells, highly aneuploid cells, and cells from adult cancers exhibit more protein buffering on average and that the degree of aneuploidy and the WGD status are potential confounding factors that need to be controlled for when analyzing average cell sample buffering.

**Figure 2:**
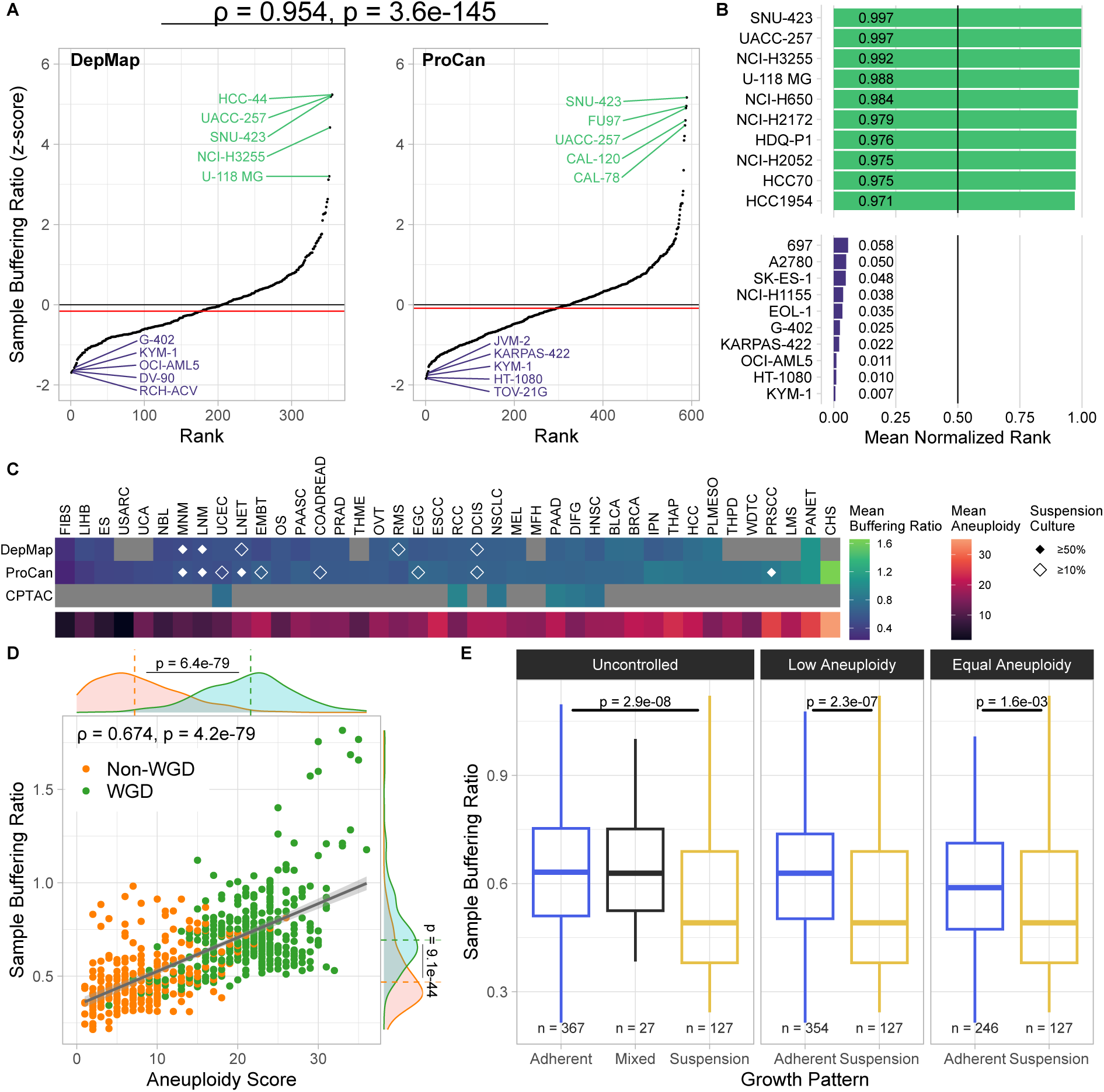
Average buffering per sample relates to cellular and clinical features of cancer. **(A)** Z-score of average buffering ratio per sample derived from pan-cancer cell line datasets (DepMap, ProCan). The sample buffering ratio z-scores between samples in DepMap and ProCan are highly correlated (Spearman’s *ρ* = 0.954*, p* = 3.6 ∗ 10*^−^*^145^). **(B)** Mean normalized ranks (MNR) of sample buffering ratios of cell lines in ProCan and DepMap. **(C)** Mean sample buffering ratio and aneuploidy score of cancer types (OncoTree code) in pan-cancer datasets. Diamond glyphs indicate the fraction of cell lines grown in suspension cultures. **(D)** Scatter plot showing a positive correlation between sample buffering ratio and aneuploidy score in the ProCan dataset (Spearman’s *ρ* = 0.674*, p* = 4.2 ∗ 10*^−^*^79^). Marginal distributions are separated and compared by whole genome doubling (WGD) status (Wilcoxon rank-sum test). **(E)** Difference in sample buffering ratio between cell lines grown in suspension and adherent cell culture types (ProCan). Controlled for aneuploidy by removing cell lines from the adherent cohort with aneuploidy scores above the maximum of the suspension cohort (middle), and by using stratified sampling to ensure an equal distribution of aneuploidy scores (right).

We noticed that non-solid cancers like myeloid and lymphoid neoplasms (MNM, LNM) and other cell lines grown in suspension cultures show significantly lower average protein buffering than those with adherent growth patterns (Figure 2E). Since aneuploidy is a confounding factor of the sample BR, and suspension-grown cell lines frequently show low AS, we controlled for differences in the degree of aneuploidy between these two growth pattern cohorts by removing cell lines from the adherent cohort with a higher aneuploidy score than cell lines grown in suspension. We additionally applied a stratified sampling scheme to the adherent growth cohort to ensure equal distributions of the aneuploidy score between the two growth pattern cohorts. Even in this setting, we observed decreased average buffering in cell lines grown in suspension cultures compared to those with adherent growth patterns (Figure 2E, Figure S2F&G). Thus, the average degree of protein buffering differs between cells and cancer types and may be an intrinsic feature affected by cellular physiology and tissue type.

### Gene dependency, protein complex participation, and random allelic expression are key predictive factors of protein buffering

To investigate which factors influence protein buffering, we compiled and integrated multiple datasets of 35 factors describing biochemical and genetic properties of each protein. We then assessed the predictive power of each factor individually to determine its effectiveness in predicting whether a protein is buffered or not by calculating the receiver operating characteristic area under the curve (ROC AUC). We calculate the ROC AUCs on BR-based (per-gene, per-sample) buffering classes derived from absolute gene copy numbers and chromosome arm copy numbers (GeneCN, ChrArm) for all pan-cancer datasets (DepMap, ProCan, CPTAC). Additionally, we determined buffering classes per gene across all samples within a dataset (ChrArm (avg.)) by a previously described method that applies average Log2FC thresholds for grouping proteins into *Scaling*, *Buffered*, and *Anti-Scaling* using chromosome arm CNAs categorized into gained, neutral, and lost chromosome arms (Schukken & Sheltzer, 2022). This resulted in three separate analysis variants (Gene CN, ChrArm, ChrArm (avg.)) providing multiple perspectives of differing granularity when analyzing factors contributing to protein buffering. We then aggregated the ROC AUC ranks of all analysis variants (Gene CN, ChrArm, ChrArm (avg.)) and all pan-cancer datasets (DepMap, ProCan, CPTAC) using mean normalized ranks (MNR), to summarize which factors are most relevant in predicting protein buffering upon gain and loss events in all samples, additionally separated by their WGD status (Supplementary Table 4).

The most important factors for predicting protein buffering upon gain were the mean dependency score (i.e., essentiality) of a gene across cell lines derived from CRISPR knock-out screens (DepMap, Broad Institute, 2023), the number of protein complexes a protein participates in (Giurgiu et al., 2019), the frequency of random allelic expression (RAE) of a gene compared to bi-allelic expression (Kravitz et al., 2023), the number of ubiquitination and methylation sites on a protein (Hornbeck et al., 2015), the probability of triplosensitivity (i.e., duplication intolerance; Collins et al., 2022) of a gene, and the number of transcription factors (TFs) that target the gene (Badia-i-Mompel et al., 2022; Garcia-Alonso et al., 2019) (Figure 3A). Upon loss, the most important factors were the mean gene dependency score, protein complex participation, the degree of non-exponential decay (NED) of a protein (Δ-score, McShane et al., 2016), RAE, and triplosensitivity (Figure 3B). This shows that there is only a partial overlap of the predictive power of factors between gains and losses. While the NED Δ-score and the probability of haploinsufficiency (i.e., deletion intolerance, Collins et al., 2022) were more predictive upon loss than upon gain events, the mean mRNA decay rate (Yang et al., 2003) was more important upon gain than loss.

**Figure 3:**
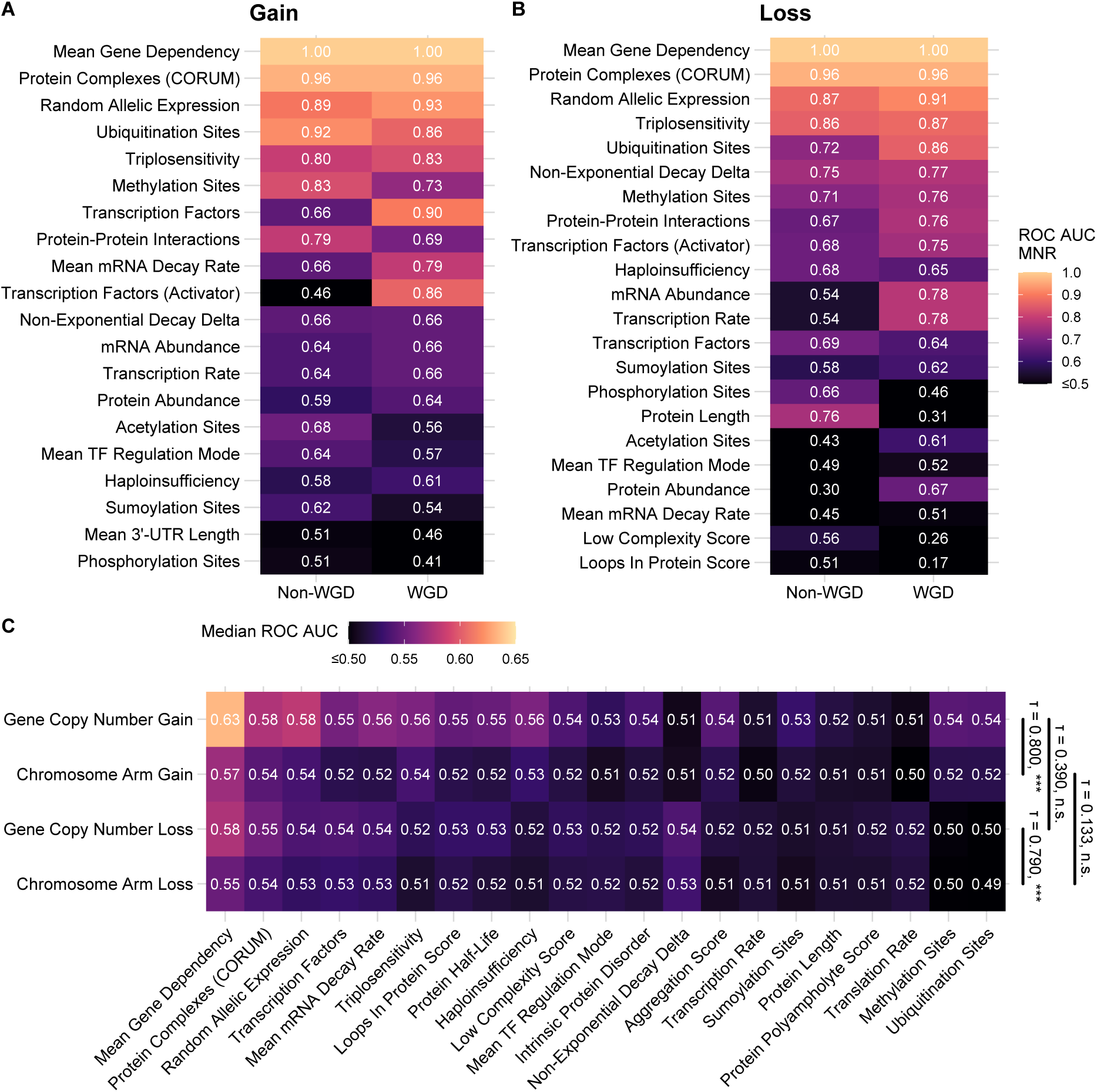
Gene dependency, protein complex participation, and random allelic expression are key predictive factors of protein buffering. **(A-B)** Mean normalized ranks (MNRs) of ROC AUCs of each factor used for classifying whether a gene is *Buffered* or *Scaling* on protein level (*Anti-Scaling* excluded). Heatmaps show a selection of factors with an MNR of at least 0.5 in one of the shown conditions. **(A)** MNRs calculated separately for whole genome doubled (WGD) and Non-WGD cell lines of the DepMap dataset across ROC AUCs of all analysis variants for gain events. **(B)** MNRs calculated separately for whole genome doubled (WGD) and Non-WGD cell lines of the DepMap dataset across ROC AUCs of all analysis variants for loss events. **(C)** Median ROC AUCs and rank correlation of factor ROC AUCs (Kendall’s *τ*) between analysis variants by bootstrapping the ProCan dataset (n = 10000). Only factors with a median ROC AUC of 0.51 across analysis variants are shown. Significance levels: (∗)*p <* 0.01, (∗∗)*p <* 0.001, (∗ ∗ ∗)*p <* 0.0001.

We next asked whether we could identify factors whose importance in predicting protein buffering is affected by genome doubling. The number of ubiquitination and methylation sites and the number of protein-protein-interactions of a protein (Alanis-Lobato et al., 2017) were more important in Non-WGD samples upon gain, while the mean mRNA decay rate (Yang et al., 2003), haploinsufficiency probability, and number of TFs with activating modes of action were more important in WGD-positive samples (Figure 3A). Comparing factors between WGD and Non-WGD samples upon loss events revealed that the number of TFs (activating + repressing), protein length, and number of phosphorylation sites were more important in Non-WGD samples. In contrast, the number of ubiquitination and acetylation sites, mRNA abundance and transcription rate (Hausser et al., 2019), and the number of protein-protein interactions were more important in WGD samples (Figure 3B).

Next, we determined whether the predictiveness of factors significantly differs between gain and loss events by bootstrapping buffering classes and calculating ROC AUC rank correlations of factors (see Methods). To control for differences caused by the choice of the analysis variant, we also evaluated the factor rank differences between buffering classes determined either by using gene copy number data or chromosome arm CNA data. The factor rank correlation was high when comparing gene copy number with chromosome arm gain (or loss) analyses, while there was no significant correlation between gain and loss analyses when using the same copy number source (Figure 3C). We therefore conclude that protein buffering is at least partially influenced by different factors upon gain than upon loss, independent of whether copy number aberration occurs on gene level or chromosome arm level.

We observed that the maximum achieved ROC AUC using the unifactorial prediction was 0.672, providing a moderate discrimination performance between *Buffered* and *Scaling* proteins (Supplementary Table 4). Furthermore, the raw values and bootstrapped ROC AUCs of factors such as protein abundance, transcription rate, and mRNA abundance were frequently and strongly pairwise correlated, indicating that these factors carry equivalent information in predicting protein buffering (Figure S3B-D). Calculating the correlation between factor value and (gene-level) Buffering Ratio confirmed mean gene dependency score, protein complex participation, triplosensitivity, mRNA decay rate, and ubiquitination sites to be important factors, with higher values being predictive for protein buffering (Figure S3E). Higher values of random allelic expression frequency and TF count were more predictive for scaling proteins. However, only a weak correlation was achieved (max(|*ρ*|) = 0.27). We conclude that a unifactorial analysis is able to identify factors with higher predictive power relative to each other, but does not enable a strong prediction of protein buffering in total.

### Explainable machine learning uncovers directional relationships between predictive factors and protein buffering

We considered that using all factors simultaneously in a combined model will improve the prediction. To approach this, we trained a series of machine learning models that use a subset of all collected factors simultaneously to predict whether a gene is *Buffered* or *Scaling*, based on the classification scheme established for the unifactorial prediction (Figure S4A). Highly correlated factors that contained redundant information were removed to prevent the models from overfitting on an arbitrary subset of factors. We derived training and test datasets from different analysis variants (Gene CN, ChrArm, ChrArm (avg.)), copy number events (gain, loss), and pan-cancer datasets (DepMap, ProCan, CPTAC), and trained the models on each training set (see Methods).

Evaluating the performance of each model on its associated test set showed that multi-factorial models perform better than using individual factors for predicting protein buffering, achieving a strong discrimination performance (Figure 4A-C; Figure S4B). Moreover, the models trained on gene copy number-derived buffering classes consistently outperformed models using chromosome arm CNA derived buffering classes. Especially upon gene copy number gain, the models showed a higher performance across datasets compared to copy number loss. This is likely because gene copy numbers provide more fine-grained copy number information, which is lost when using chromosome arm copy numbers. This information can then be used in conjunction with the BR to generate more accurate buffering classes that can be predicted more confidently. Additionally, the performance of tumor sample derived models (CPTAC, Gene CN gain: ROC AUC = 0.846, loss: ROC AUC = 0.811) surpassed the predictive performance of models trained on cell line data (DepMap, Gene CN gain: ROC AUC = 0.773, loss: ROC AUC = 0.684). Next, we evaluated the prediction performance of the models on unseen datasets (out-of-sample prediction). We observed that models trained on cell line datasets perform well on other cell line datasets upon gene copy number gain, but show a weaker performance on tumor sample data and vice versa (Figure S4C). Based on these observations we assume that factors affect protein buffering slightly differently in tumor samples and cell lines.

**Figure 4:**
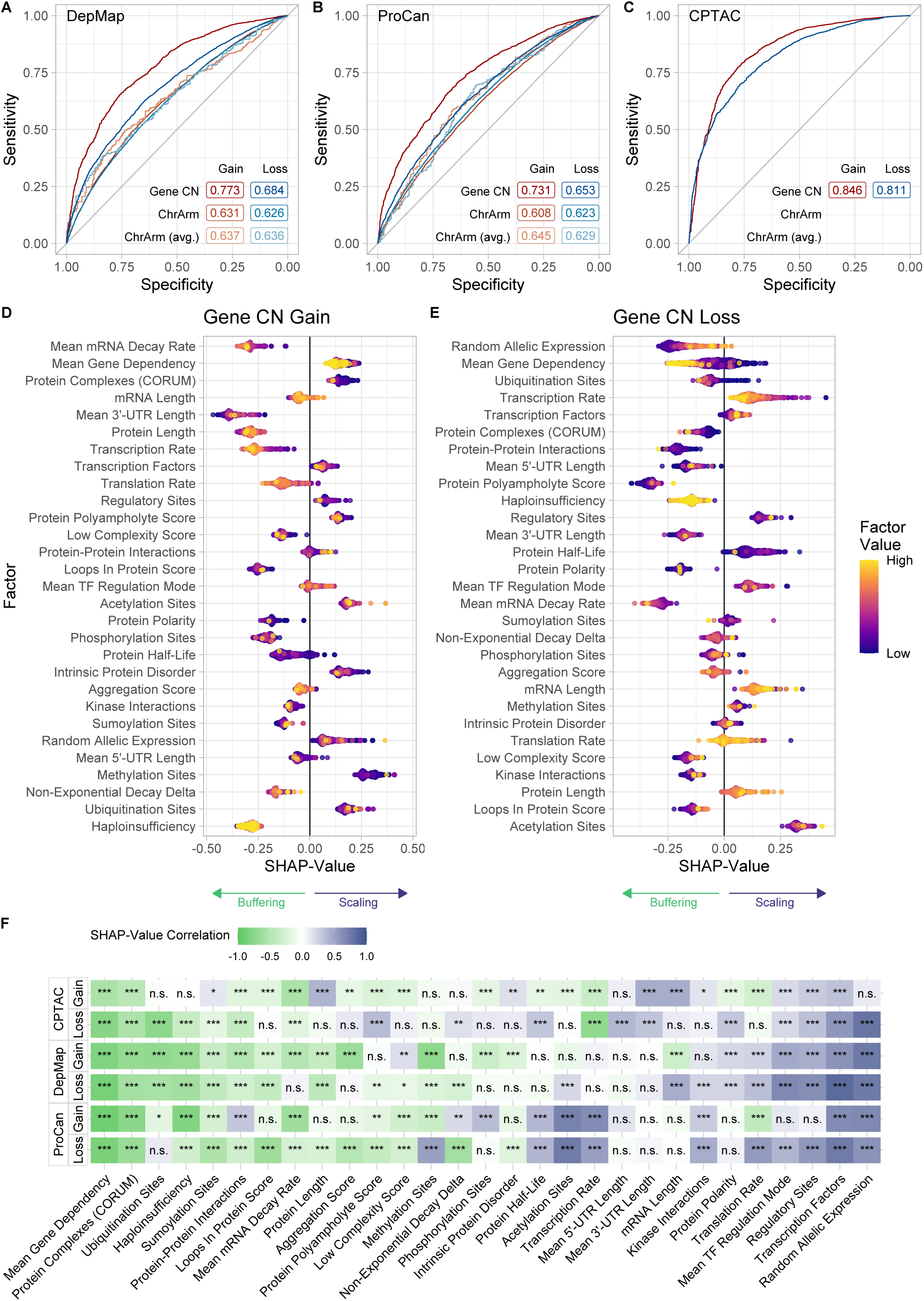
Explainable machine learning uncovers directional relationships between pre-dictive factors and protein buffering. **(A-C)** ROC curves and AUCs of multifactorial xgbLinear models trained on predicting *Buffered* and *Scaling* classes for different copy number events, analysis variants and datasets. The ROC curves depict the model’s performance on its respective test set (80/20 training/test-split). **(D, E)** SHAP values generated by evaluating the trained CPTAC gene copy number models on a random subset of the respective test set (n = 300). Negative SHAP values depict a higher contribution of a factor within the model towards predicting an observation as *Buffered*. Each dot represents a gene in a sample. Color represents the min-max-scaled value of the underlying factor used for prediction in the model for this observation. **(F)** Heatmap showing the correlation (Spearman’s *ρ*) between SHAP values estimated on a random subset of the respective test set (n = 300) and their underlying factor values used for prediction. Negative correlation indicates stronger contribution of a factor towards predicting observations as *Buffered* if the factor value is high. Correlation is shown for pan-cancer datasets (DepMap, ProCan, CPTAC) upon gene copy number gain and loss with significance levels indicated (Benjamini-Hochberg adjusted p-values, (∗)*p_adj_ <* 0.01, (∗∗)*p_adj_ <* 0.001, (∗ ∗ ∗)*p_adj_ <* 0.0001)

To determine how each factor contributed to the prediction of each model, we calculated SHapley Additive exPlanation values (SHAP values, Lundberg & Lee, 2017), which quantify the contribution of each feature to a model’s prediction by attributing the differ-ence between the actual and baseline predictions to individual features. As SHAP values can be obtained for each factor and prediction, we calculated SHAP values on a subset of the model’s test set using only observations where the model’s prediction was correct. Negative SHAP values indicated that for a given protein in a sample the model is more confident towards classifying the observation as *Buffered* using the corresponding factor value. For example, higher values of mean gene dependency were associated with lower SHAP values, meaning that the model was more confident in predicting a protein in a sample as *Buffered* if the gene is essential in many samples (Figure 4D&E).

To quantify the direction of influence of a factor in predicting protein buffering, we calculated the Spearman correlation between SHAP values and corresponding factor values for each factor. This approach achieved a stronger correlation than by directly correlating BRs and factor values as done in the unifactorial analysis (max(|*ρ*|) = 0.95). We observed that frequent participation in macromolecular complexes and a high mean gene dependency are consistently predictive for *Buffered* proteins upon gene copy number gain and loss (Figure 4F). In contrast, a high number of TFs binding to a gene, high fraction of TFs with activating regulatory modes, high RAE frequency, and high number of sites known for regulating a protein’s function are consistently predictive for *Scaling* proteins upon gene copy number gain and loss. Interestingly, an increased number of ubiquitination sites is more predictive for *Buffered* proteins upon gain than upon loss in DepMap and ProCan, whereas in CPTAC an increased number of ubiquitination sites is more predictive for *Buffered* proteins upon gene copy number loss. We also observed this pattern in the unifactorial correlation analysis (Figure S3E).

In conclusion, multifactorial machine learning models outperform unifactorial analyses in predicting protein buffering. We showed that increased mean gene dependency and protein complex participation contribute to a higher probability of protein buffering, whereas a higher number of activating TF interactions, protein regulatory sites, and higher RAE frequency contribute to a lower probability of protein buffering.

### Cancer samples with high average buffering show altered differential expression patterns related to protein folding

Next, we investigated if high average protein buffering in cell lines and tumor samples is associated with specific gene expression patterns. Using the previously calculated sample BRs, we split the samples into low- and high-buffering groups, and inspected which proteins were differentially expressed in high-buffering samples compared to low buffering samples (Figure 5A). We identified the cancer-drivers LMNA (structural protein of the nuclear lamina lamin A), EGFR (transmembrane tyrosine kinase receptor of the epidermal growth factor), RRAS (member of the RAS family of small GTPases), CTNNB1 (*β*-catenin, a key component of the Wnt signaling pathway), and many others, as significantly upregulated in high-buffering cell lines in both pan-cancer cell line datasets (DepMap, ProCan). Among proteins downregulated in high-buffering cell lines in ProCan we observed LCP1 (an actin binding protein linked to hereditary cancer syndrome) and DNA methyltransferase DNMT1. ORA of the downregulated genes in cell lines with high average buffering revealed that terms related to protein folding, such as the chaperonin containing complex CCT, DNA replication, and nucleotide binding were enriched (Fig-ure 5B). The CCT/TRiC complex is involved in folding 10% of the proteome including actins, tubulins, and cell cycle regulators (Gestaut et al., 2019). Among the set of up-regulated proteins, integrin complex partners, cell adhesion proteins, cadherin binding proteins, and proteins involved in the organization of the extracellular matrix were en-riched (Figure 5C).

**Figure 5:**
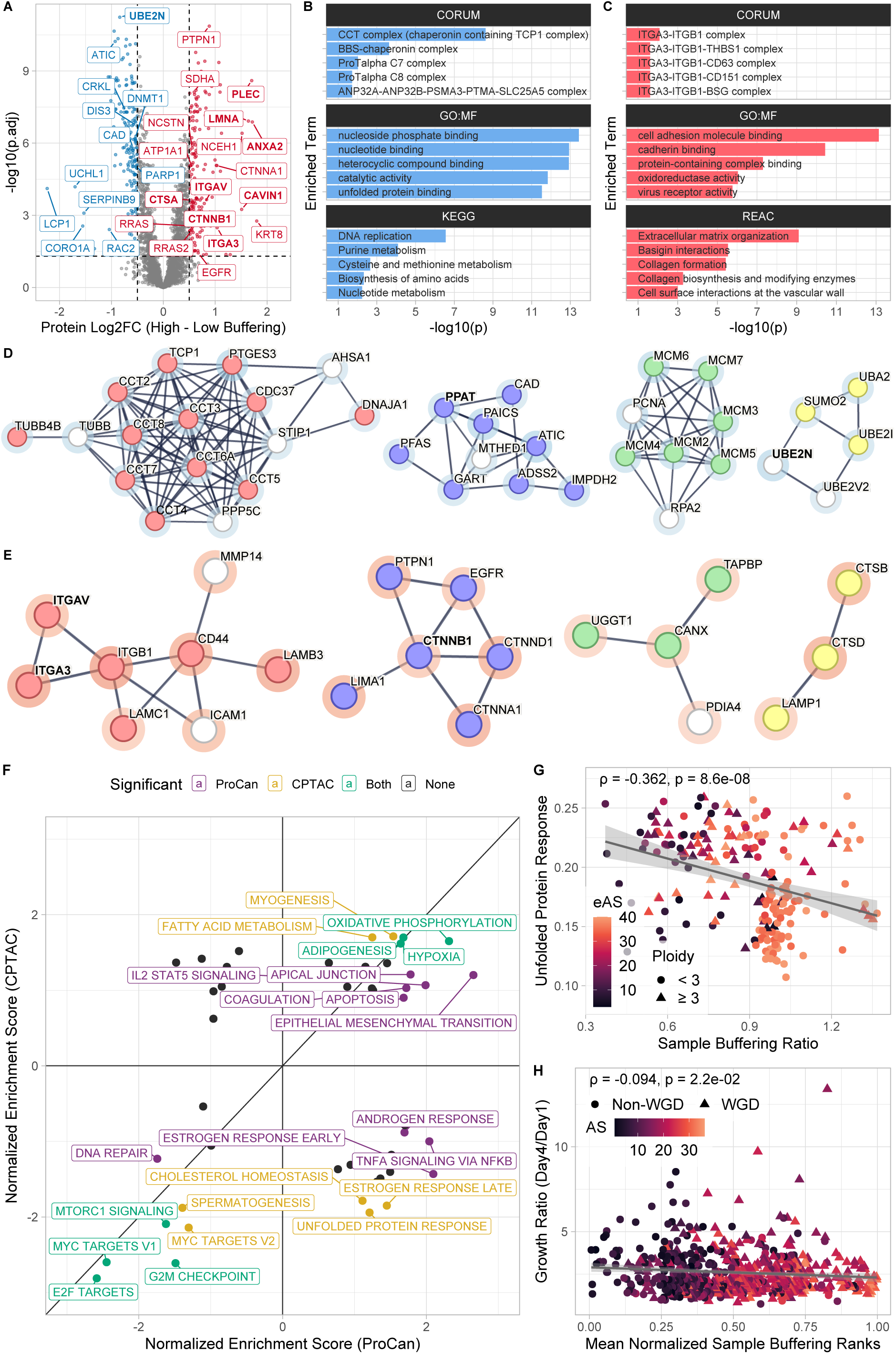
Cancer samples with high average buffering show altered differential expression patterns related to protein folding. **(A)** Volcano plot showing a log2 fold-change between high (>80% quantile of sample buffering ratios) and low (<20% quantile) buffering cell lines in the ProCan dataset plotted against the Benjamini-Hochberg adjusted student’s t-test p-values. Labeled are genes with significant differential expression (|*Log*2*F C*| *>* 0.5*, p_adj_ <* 0.05). Genes with bold labels are significant in all cell line and control datasets. **(B, C)** Over-representation analysis (ORA) of genes that are significantly down-**(B)** and up-regulated **(C)** in highly buffering cell lines. **(D)** Selected network clusters within the set of significantly down-regulated genes in highly buffering cell lines. Highlighted gene sets: *unfolded protein binding* (red, GO:MF), *nucleotide biosynthesis* (blue, Reactome), *MCM complex* (green, GO:CC), and *protein sumoylation* (yellow, GO:BP). **(E)** Selected network clusters within the set of significantly up-regulated genes in highly buffering cell lines. Highlighted gene sets: *ECM receptor interaction* (red, KEGG), *Cadherin binding* (blue, GO:MF), *unfolded protein binding* (green, GO:MF), and *lysosome* (yellow, KEGG). The rings around the dots indicate the log2-fold change (blue: negative, red: positive). **(F)** 2D enrichment plot comparing the normalized enrichment scores of MSigDB HALLMARK gene sets between CPTAC and ProCan. Significance cutoff of Benjamini-Hochberg adjusted p-values: *p_adj_ <* 0.05. **(G)** Scatter plot showing a negative correlation between sample buffering ratios and single-sample enrichment scores for the *Unfolded Protein Response* HALLMARK gene set using CPTAC tumor sample proteome data (Spearman’s *ρ* = −0.362*, p* = 8.6 ∗ 10*^−^*^08^). Color indicates the estimated aneuploidy score of a tumor sample. **(H)** Scatter plot showing a weak negative correlation between mean normalized buffering ratio ranks and the proliferation (day1-to-day4 growth ratio) of corresponding cell lines (Spearman’s *ρ* = −0.094*, p* = 0.022). Color represents a cell line’s aneuploidy score.

We noticed that blood cancers, which are frequently grown in suspension, have particularly low sample BRs. To avoid confounding effects of the cell growth method, we created an adherent control by calculating the differential expression between high and low buffering cell lines that were only grown adherently. Using this adherent control, cell adhesion, cell motility, and extracellular matrix organizing proteins were still upregulated, while nucleotide metabolism and biosynthesis proteins remained downregulated in high-buffering cells (Figure S5A-C). We furthermore found 27 genes that were commonly deregulated upon high buffering in all cell line and adherent control datasets (Supplementary Table 8).

STRING was used to visualize how differentially expressed genes in ProCan are involved in known and predicted protein-protein interactions (Szklarczyk et al., 2023; Figure 5D&E). In addition to the CCT/TRiC complex, we observed that genes involved in nucleotide biosynthesis, protein sumoylation-related genes, and DNA replication factors associated with the MCM complex were downregulated (Figure 5D). The deregulation of the MCM complex has been linked to increased genomic instability and to aneuploidy (Bochman & Schwacha, 2009; Passerini et al., 2016; Shima et al., 2007). Furthermore, both PCNA and UBE2N were down-regulated; these genes play a role in error-free post-replication repair (Hoege et al., 2002; S. Kim et al., 2024; Moldovan et al., 2007; Motegi et al., 2008; Unk et al., 2008). In agreement with the ORA results, clusters containing integrins and proteins related to extra-cellular matrix (ECM) receptor interaction were upregulated (Figure 5E). In particular, the integrins ITGA3, ITGA5, and ITGB1 were upregulated in highly buffering cell lines; their deregulation has been shown to alter cell adhesion and promote cell invasion (Fukushi et al., 2004; Mueller et al., 1999). Cadherin-binding proteins were upregulated as well, including the oncogene EGFR which positively regulates cell migration (Ghosh et al., 2010), and Catenin beta-1 (CTNNB1) involved in the regulation of cell adhesion (Brembeck et al., 2006; W. K. Kim et al., 2019). We observed that EGFR, among other oncogenes, had a higher BR upon chromosome arm loss, implying that the upregulation of EGFR in highly buffering cells is due to EGFR itself being buffered upon loss (Figure S5E-G). EGFR has been shown to frequently scale with chromosome arm gain (Schukken & Sheltzer, 2022), which has been confirmed in our analysis (Fisher’s Exact test, *OR* = 0.27*, p* = 7.9 ∗ 10*^−^*^4^). This suggests that cancer cell lines retain elevated EGFR levels independent of chromosome arm gain or loss. Furthermore, the lysosomal proteins Lysosomal-membrane-associated glycoprotein 1 (LAMP1), Cathepsin D (CTSD), and Cathepsin B (CTSB) were upregulated. CTSD and CTSB overexpression as well as increased LAMP1 cell-surface expression is associated with increased metastatic potential and tumor invasion (Jensen et al., 2013; Lee et al., 2024; Rochefort et al., 1990; Ruan et al., 2016; Sarafian et al., 1998). Similar to the downregulated genes, we also found clusters of proteins involved in unfolded protein binding in upregulated genes (UGGT1, CANX, TAPBP), playing a role in the retention and quality control of unfolded and misfolded proteins in the endoplasmic reticulum (Adams et al., 2020; Swanton et al., 2003). This in combination with the downregulation of the CCT/TRiC complex suggests an increase in protein folding quality control in cell lines with high average protein buffering while demand for overall protein folding may be decreased.

To further investigate which cancer hallmarks are deregulated in cells with high average buffering, we applied gene set enrichment analysis (GSEA) using the MSigDB hallmark gene sets and compared enrichment scores between tumor samples and cell lines. MYC target, G2M checkpoint, MTORC1 signaling and E2F target gene sets were significantly downregulated in both high-buffering tumor samples and cell lines, while DNA repair gene sets were significantly downregulated in cell lines (Figure 5F). As we identified aneuploidy as a potential confounding variable for BR-based analyses, we also performed GSEA on differential expression data between high and low aneuploid samples and compared the results to the GSEA analysis results between high and low buffering samples. DNA repair, E2F target, and MTORC1 signaling pathways were upregulated in highly aneuploid tumor samples, showing that the downregulation of these pathways were unique to tumor samples with high average buffering and independent of the influence of aneuploidy (Figure S5H).

Highly aneuploid cell lines and primary tumors suffer from increased proteotoxic stress and have an increased Unfolded Protein Response (UPR) (Ippolito et al., 2024; Oromendia & Amon, 2014). We observed that the UPR gene set was significantly downregulated in tumor samples with high average buffering, but not in cell lines, suggesting that tumor samples could use protein buffering to alleviate proteotoxic stress and do so more effectively than cell lines (Figure 5F). To test this hypothesis, we applied single-sample GSEA (ssGSEA, Barbie et al., 2009) to CPTAC using the UPR hallmark gene set and correlated the single-sample enrichment scores with the sample BR. Indeed, tumor samples that exhibit high average buffering showed a decreased UPR, especially in highly aneuploid WGD-negative samples, implying that protein buffering reduces proteotoxic stress (Figure 5G). Sub-tumor heterogeneity and low tumor purity could potentially confound these results, but the UPR enrichment score and tumor purity were not significantly correlated (Spearman’s *ρ* = −0.025*, p* = 0.72; Figure S5) and tumors with a purity above 70% exhibited an increased UPR (UPR > 0.2) less frequently than low-purity tumors (*OR* = 0.41*, p* = 0.021). Futhermore, the correlation between sample BR and UPR enrichment score increased when removing tumor samples with a purity below 50% (Spearman’s *ρ* = −0.43*, p* = 6.4 ∗ 10*^−^*^3^), suggesting that the results are not confounded by tumor samples with low purity.

Therefore, we asked whether cell lines exhibiting high levels of protein buffering could have a proliferative advantage, as they might be able to better mitigate the proteotoxic stress introduced by aneuploidy. Using growth rates from DMSO-treated cell lines in the GDSC drug screen dataset (Iorio et al., 2016) showed that some cell lines proliferated slightly worse the more aneuploid they were (particularly WGD-positive cell lines: Spearman’s *ρ* = −0.270*, p* = 8.2 ∗ 10*^−^*^6^). However, we did not observe a strongly significant correlation between growth rates and average buffering estimates in cell lines (Figure 5H). We conclude that high buffering alters protein homeostasis in cell lines, affecting DNA repair, protein folding, and cell adhesion pathways. Interestingly, increased buffering did not provide a general proliferative advantage.

### CRISPR knock-out screens reveal vulnerabilities of weakly buf-fering cells

As high average protein buffering does not appear to provide a general proliferative advantage in aneuploid cell lines, we investigated whether these cell lines show differential dependency on certain genes or if increased buffering mitigates aneuploidy-induced vul-nerabilities. We divided the cell lines into groups with high and low average buffering (>80%, <20% sample BR MNR), and compared the dependency scores of genes between these groups. The dependency scores were derived from CRISPR-KO screens and quantify how dependent a cell line is on a particular gene. Cell lines with high average buffering were more dependent on the oncogenes CRKL (a protein kinase possibly activating RAS and JUN) and Hippo signaling pathway transcription coactivator WWTR1. On the other hand, they are less dependent on the oncogenes MDM4 (a p53 regulator), TAF1 (TATA binding factor); tumor suppressor genes POT1 (telomere-protecting protein), SDHD (sub-unit of the mitochondrial respiratory chain), and a sensor of oxidative stress KEAP1 (Figure 6A). Furthermore, ITGAV, FERMT2, PTK2, GPR61 and RELA were exclus-ively essential in high-buffering cells, while CENPI, DROSHA, RNF40, TUBD1, TMX2, RAD1, DPY30, PTBP1, TBCE, MED10, WDR48, TUBE1, and NAMPT were exclus-ively essential in low buffering cell lines.

**Figure 6:**
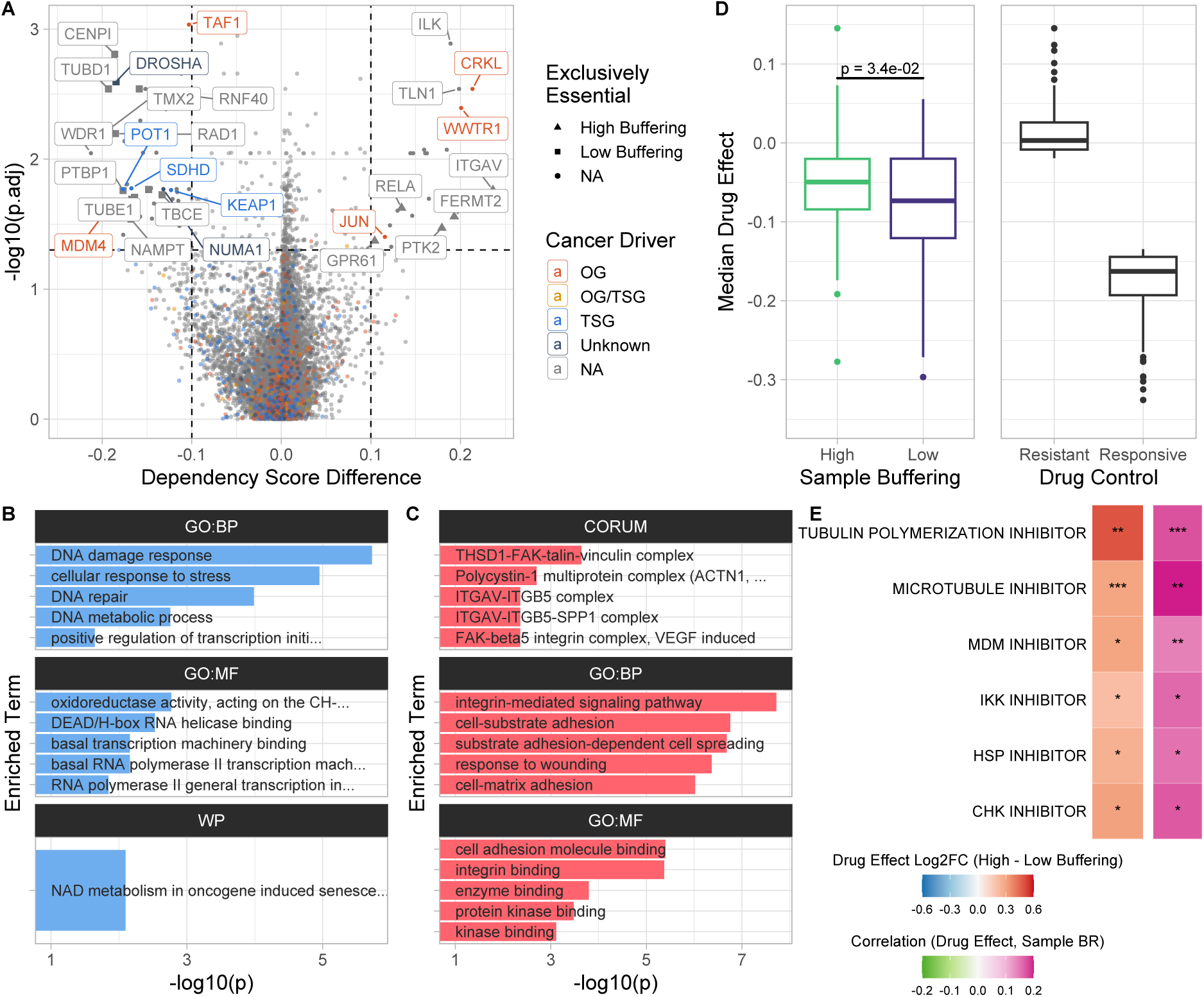
CRISPR knock-out screens reveal vulnerabilities of weakly buffering cells. **(A)** Volcano plot showing the difference of the CRISPR-KO dependency score between high (>80% quantile of sample BR MNRs) and low (<20% quantile) buffering cell lines plotted against Benjamini-Hochberg adjusted Wilcoxon rank-sum test p-values. **(B, C)** Over-representation analysis (ORA) of genes with significantly reduced **(B)** and increased **(C)** CRISPR-KO dependency scores in highly buffering cell lines. **(D)** Median drug effect on cell viability across drugs from the PRISM Repurposing dataset for high (>80% quantile of sample BR MNRs) and low (<20% quantile) buffering cell lines. Cell lines resistant (>80% quantile of median drug effect) and responsive (<20% quantile) to drug treatment are shown as controls. **(E)** Drug mechanisms with significant log2 fold-change of mean drug effect between high and low buffering cell lines (Student’s t-test, |*Log*2*F C*| *>* 0.2*, p_adj_ <* 0.05) and significant correlation between sample BR MNRs and drug effect (Spearman’s *ρ*) with significance levels shown (Benjamini-Hochberg adjusted p-values, (∗)*p_adj_ <* 0.05, (∗∗)*p_adj_ <* 0.01, (∗ ∗ ∗)*p_adj_ <* 0.001).

ORA of the genes with significant dependency score differences showed that cell lines with high average buffering were less dependent on DNA damage response, DNA repair, and cellular stress response genes than cell lines with low average buffering (Figure 6B). Interestingly, high-buffering cells were more dependent on cell adhesion regulation, substrate adhesion-dependent cell spreading, kinase binding, and integrin binding genes (Figure 6C). As before we created a control subset of cell lines that were adherently grown, and compared the differential dependency between high and low buffering cell lines within this subset (Figure S6A, Supplementary Table 9). ORA showed that general transcription initiation factor binding genes were less essential in high-buffering cell lines (Figure S6B). Furthermore, genes that remained exclusively essential in low buffering cell lines after controlling for growth method were CENPI (Centromere Protein I), RAD1 (RAD1 Checkpoint DNA Exonuclease), TUBD1 (Tubulin Delta 1), and TBCE (tubulin folding protein). This analysis further highlights the physiological differences between high and low buffering cell lines.

Next, we analyzed whether samples with different degrees of average protein buffering exhibit an altered sensitivity to drug treatment. We collected drug effect scores quantifying the changes in cell viability between drug and DMSO treatment from the PRISM Repurposing dataset (Corsello et al., 2020; DepMap, Broad Institute, 2023), where lower values indicate a decreased growth of the cells after drug treatment compared to the control using DMSO. We categorized cell lines as drug-resistant if their median drug effect was higher than 80% of the drug effect scores, while cell lines with a median drug effect score below the 20% percentile were categorized as drug responsive. As before, we also categorized cell lines as high and low buffering based on the 80% and 20% percentile of the sample BR MNR. Remarkably, the median drug effect on cell viability of high-buffering cells is closer to the drug-resistant cell cohort, and the drugs have a significantly lower median effect on the viability of high-buffering cells compared to cell lines with low average protein buffering, although this difference is small (Figure 6D). To test whether high buffering and drug resistance are frequently co-occurring, we separated all available cell lines by median sample BR MNR and median drug effect. This showed that high-buffering cell lines were more frequently drug-resistant than cell lines with a sample BR MNR below median (Fisher’s exact test, *OR* = 1.525*, p* = 2.1 ∗ 10*^−^*^2^). The increased drug resistance was largely explained by increased aneuploidy, however, high buffering samples were still slightly more frequently drug resistant independent of aneuploidy status (Figure S6C-E; High Aneuploidy: *OR* = 1.088*, p* = 0.021; Low Aneuploidy: *OR* = 1.182*, p* = 0.57). While highly buffering cell lines showed no sensitivity to any group of drugs, the low buffering cell lines were significantly more sensitive to HSP, CHK, MDM, IKK, microtubule, and tubulin polymerization inhibitors (Figure 6E). Thus, cells that cannot efficiently buffer protein imbalance arising from aneuploidy are particularly sensitive to drugs that target protein folding, mitotic spindle function, innate immune response, and DNA damage response. As a control, we analyzed the drug effect difference between high and low aneuploid cells. Drug mechanisms that were not significant in highly aneuploid cells and were unique to low buffering cells were CHK and HSP inhibitors (Supplementary Table 10). In conclusion, efficiently buffering aneuploid cells showed increased resistance to drugs in general and did not respond to any of the tested drug mechanisms, indicating that these cells are particularly difficult to treat.

## Discussion

Abnormal chromosomal content, or aneuploidy, strongly alters protein homeostasis due to aberrant expression of genes located on the affected chromosomes. Yet, a significant fraction of the aberrant expression caused by altered gene dosage is mitigated by adjusting the protein abundance (Chunduri et al., 2021; Dephoure et al., 2014; Stingele et al., 2012). This means protein abundance in aneuploid cells often resembles diploid levels rather than the abundance expected based on the gene dosage.

Previous research indicated that gene dosage buffering is widespread in aneuploid cell lines and that its degree is affected by certain gene and protein features, such as the number of ubiquitylation sites, or participation in macromolecular complexes (Schukken & Sheltzer, 2022). To obtain additional insight into the factors affecting dosage compensation, we established a novel indicator we termed “Buffering Ratio” (BR), where we quantified the protein-level attenuation for each gene within single samples. This approach was designed to deliver a gene copy number-sensitive quantification of protein abundance changes in aneuploid samples that may be “averaged out” when using protein-average Log2FC cutoffs as in Schukken & Sheltzer (Figure 1B). While this approach allows us to evaluate the effect of whole-genome doubling, or identify variability of buffering, it also has its limitations. In particular, the BR is sensitive to noise from both proteomics and copy number data. To circumvent these issues, we calculated confidence scores for the BRs, and removed BR measurements with low confidence.

Using this approach, we demonstrate that gene dosage compensation occurs not only in aneuploid *in vitro* cancer cell lines but also extensively and efficiently in aneuploid *in vivo* tumor samples (Figure 1). The widespread phenomenon of gene dosage buffering highlights the importance of maintaining protein homeostasis despite aberrant chromosome and gene copy numbers. While buffering in pan-cancer cell lines and tumor samples is comparable, chromosome-engineered aneuploid human cell lines are buffered to a greater extent. The underlying causes of these differences remain unclear. We hypothesize that the distinct genomic alterations in each sample type may play a role: engineered cell lines exhibit gains and losses of individual chromosomes, whereas cancer samples typically harbor copy number variations affecting multiple chromosomes, as well as plethora of mutations.

To identify factors that contribute to gene dosage buffering, we applied both unifactorial and multifactorial models to analyze their effects and predictive capacity. While uni-factorial models predict factors contributing to buffering with a low predictive power (∼0.6 ROC AUC, Figure 3; Schukken & Sheltzer, 2022), our novel multifactorial ML models performed significantly better, with up to 0.85 ROC AUC (Figure 4). Using these models, we established gene dependency as the key factor in gene dosage buffering after chromosome or gene gains and losses in all analyzed datasets. This demonstrates that the abundance of essential genes must be maintained at optimal levels independently of the correspond-ing gene copy number. Additionally, being a member of macromolecular complexes, as well as having a high number of ubiquitylated sites is strongly predictive of buffering, as previously observed in other works (Figure 3 & 4; Ishikawa et al., 2017; Schukken & Sheltzer, 2022; Stingele et al., 2012). This supports the hypothesis that protein buffering is in parts conveyed by degradation of non-stochiometric subunits of macromolecular complexes. In contrast, scaling was affected mainly by transcriptional features (transcription factors, bi-allelic expression, etc.) and mRNA turnover. Remarkably, we found a different predictive power of individual factors for buffering protein abundance upon gene/chromosome gain and loss (Figure 3). This suggests that chromosome gains and losses may affect cellular physiology differently and mitigation of the protein imbalance occurs partly by different mechanisms. Out-of-sample prediction showed that our models performed well across datasets, highlighting the reproducibility of our results. However, the predictiveness differed between ML models trained on cell line datasets compared to tumor sample-trained models. This suggests, that tumor samples buffer changes in protein abundance to a different extent than cell lines do.

Our approach of buffering ratio quantification also allowed us to differentiate the consequences of gene copy number changes among cell lines and tumor types. Some cell lines were extremely effective in buffering, while others did not compensate for the changes in copy number as much (Figure 2). Tumor samples also showed this variability. Cell lines grown in suspension showed lower levels of average buffering than cell lines with adherent growth patterns. Interestingly, our analysis suggests that aneuploidy is a strong confounding factor of gene dosage buffering, as we observed a strong correlation of the BR with the respective aneuploidy score (AS) of each cell line. Separating aneuploidy and buffering is difficult, as only aneuploid cells present a protein imbalance that can be buffered. The fact that highly aneuploid cells buffer the gene dosage more efficiently than cells with low AS may also suggest that aneuploid cells adaptively rewire their gene expression network to minimize the impact of gene dosage changes. Other possible confounding factors (e.g., p53 status) showed negligible effects. We hypothesize that relatively low levels of buffering in pediatric cancers suggest that these cancers developed quickly and have not adapted to altered protein homeostasis. In contrast, the cancer types with increased buffering suggest an etiology related to environmental and age-related factors. These cells might have gained different aneuploidies in multiple steps and adapted to aneuploid-induced stresses through buffering, leading to a more balanced proteostasis.

We found that cell lines with highly efficient gene dosage buffering downregulated the chaperonin complex CCT/TRiC, replication factors, and factors required for nucleotide biosynthesis. In contrast, pathways related to integrin signaling, extracellular matrix, and cell adhesion were upregulated in highly buffering aneuploid cells, even when controlling for confounding factors. Our analyses point out that protein folding plays a critical role in buffering, and that buffering correlates with DNA and RNA metabolism, as well as with extracellular matrix regulation (Figure 5). But which of these consequences are due to the efficient buffering, and what is a cellular response to aneuploidy? To disentangle these interlinked phenomena, we performed gene set enrichment analysis (GSEA) for high vs. low buffering samples and compared the results with analysis in high vs. low aneuploid samples. This approach clearly shows that DNA repair, E2F targets, and mTORC1 targets are differentially regulated in samples with high buffering ratio, independently of aneuploidy. Tumor samples with higher average buffering also showed a reduced unfolded protein response, particularly in WGD-negative, but highly aneuploid cells. Controlling for confounding effects of tumor heterogeneity confirmed these results, indicating that buffering alleviates proteotoxic stress introduced by aneuploidy. Furthermore, the upregulation of oncogenes in high-buffering cells suggests that buffering affects the oncogenic potential of cells, in particular by increasing the abundance EGFR upon loss of chromosome arm 7p (Figure 5A, S5E-G).

The differences in buffering capacity between tumor samples and cell lines, as well as between cancer types, raised the question of whether efficient buffering influences their growth, survival, and response to treatment. This was suggested by recent observations that naturally occurring adapted aneuploid strains of *Saccharomyces cerevisiae* efficiently buffer the gene dosage by increasing their ubiquitin-proteasome-dependent protein turnover (Muenzner et al., 2024). Intriguingly, we observed no direct improvement of proliferation in efficiently buffering cell lines compared to low-buffering cells. In fact, highly aneuploid cancer cell lines showed a decreased proliferation compared to low aneuploid cells *in-vitro*. Furthermore, we observed that increased buffering is associated with high aneuploidy. We, therefore, propose that buffering mitigates the fitness costs of aneuploidy and thus enables increased tolerance to aneuploidy. It should be noted that this analysis has been performed only for cell lines in a pan-cancer analysis that does not control for genetic and tissue-specific differences. The conditions may be different from *in vivo* tumor growth. However, There are no data currently available which would allow to directly test the hypothesis that increased buffering is an adaptive feature and facilitates the proliferation of aneuploid tumors.

Finally, we considered the possibility that the buffering capacity may affect gene dependency or drug sensitivity. Strikingly, we found that high-buffering cells exhibit increased drug resistance and are less dependent on genes related to DNA damage repair and response to cellular stresses, but more dependent on genes involved in cell adhesion. This pattern aligns with their differential expression profiles (Figure 5 & 6), but the surprising link between cell adhesion and gene dosage buffering remains unclear. We note that increased aneuploidy is likely the dominating cause of increased drug resistance in high-buffering cells (Andrade et al., 2023). In contrast, low-buffering cells are generally more sensitive, in particular to HSP, MDM, CHK, IKK, microtubule, and tubulin polymerization inhibitors. Intriguingly, low-buffering cells remained sensitive to HSP and CHK inhibitors even when controlling for aneuploidy. These findings support the hypothesis that low-buffering cells experience greater proteotoxic stress and that inhibiting chaperones in these cells further exacerbate these stresses, thereby reducing growth. Furthermore, TUBD1 (Tubulin Delta 1), TBCE (tubulin folding protein), and CENPI (Centromere Protein I) participate in the correct formation of the mitotic spindle and are essential in low buffering cells, suggesting that cells with weakly buffered protein abundance rely on the proper progression of mitosis, as they cannot tolerate increased aneuploidy.

Based on our results we hypothesize that cells that efficiently buffer changes in protein abundance have an adaptive advantage by altering the adhesion to the extracellular matrix, and alleviating aneuploidy-induced proteotoxic stress. Future research should uncover the links between gene dosage buffering and further factors identified in our analysis.

## Methods

### Cell Line Generation

p53 deficient hTert immortalized human retinal pigment epithelium (RPE1p53-/-; Mardin et al., 2015) cell lines were cultured in DMEM supplemented with GlutaMAX™, 10% fetal bovine serum (FBS; Gibco), and 1% Penicillin-Streptomycin (Gibco). All cell lines were cultured at 37 *^◦^*C in a humidified incubator with 5% CO_2_. To generate aneuploid cell lines we used an adapted protocol from Soto et al. based on a combination of CENP-E and MPS1 inhibition (Soto et al., 2017). Stable cell lines were grown and sequenced to determine the karyotype. The obtained aneuploid cell lines were labeled RM13 (monosomy 13) and Rtr13 (trisomy 13). Cell lines were routinely checked for mycoplasma contamination (MycoStrip detection kit by InvivoGen) and authenticated through sequencing.

### MS Sample Preparation and Labeling of Peptides by Tandem Mass Tags (TMT)

Peptide preparation and TMT labeling were carried out following the manufacturer’s protocol. Briefly, cells were harvested by trypsinization from plates. 10^6^ cells were washed twice with PBS, pelleted, and stored at −80 *^◦^*C. Cells were lysed by adding 100 μL lysis buffer (10% SDS in 100 mM Triethylammonium bicarbonate (TEAB)) using strong ultrasonication. The lysates were clarified via centrifugation at 16, 000 × *g* for 10 minutes at 4 *^◦^*C, and protein concentrations were determined using the BCA protein assay kit (Thermo Scientific). 50 μg of protein were reduced using 5 mM Tris 2-carboxyethylphosphine (TCEP) for 1 hour at 55 *^◦^*C and alkylated with 10 mM iodoacetamide in the dark for 30 minutes at 25 *^◦^*C. The proteins were then precipitated overnight at −20 *^◦^*C by adding six volumes of acetone. After precipitation, the proteins were resuspended in 100 mM TEAB (pH 8.5) and digested overnight at 37 *^◦^*C with sequencing-grade modified trypsin. For TMT labeling, the trypsin-digested peptide samples were labeled with isobaric tags (TMT 10-plex, Thermo Fisher Scientific), with individual tags assigned as follows: RM13_01 (TMT126), RM13_02 (TMT127N), RM13_03 (TMT127C), RPE1_p53-/-_01 (TMT128N), RPE1_p53-/-_01 (TMT1228C), RPE1_p53-/-_03 (TMT129N), Rtr13_01 (TMT129C), Rtr13_02 (TMT130N), Rtr13_03 (TMT130C), and equal mixture of all cell samples (TMT131). After labeling the reaction was quenched by adding 5% hydroxylamine and incubation for 15 min at room temperature. Then all samples were combined in equal amounts and dried under vacuum. Pooled peptides were fractionated into eight fractions (5, 10, 12.5, 15, 17.5, 20, 22.5, 25, 50% ACN) using the Pierce^™^ High pH Reversed Phase Peptide Fractionation Kit (Thermo Scientific). Each fraction was dried under vacuum, resuspended in buffer A (0.1% formic acid) containing 0.01% trifluoro acetic acid and analyzed using nanoflow liquid chromatography coupled to a Q-Exactive HF mass spectrometer (Thermo Fisher Scientific). Peptide separation was carried out with an EASY-nLC 1200 ultra-high-pressure liquid chromatography system using chromatography columns (50 cm, 75 mm inner diameter) packed in house with ReproSil-Pur C18-AQ 1.9 μm resin (Dr. Maisch GmbH). Peptides were loaded in buffer A (0.1% formic acid) and eluted with a non-linear gradient of 5-60% buffer B (0.1% formic acid, 80% acetonitrile) at a rate of 250 nL*/*min over 180 min. Column temperature was maintained at 60 *^◦^*C. Data acquisition alternated between a full scan (120 K resolution, maximum injection time of 8 ms, AGC target of 3e6) and 15 data-dependent MS/MS scans (60 K resolution, maximum injection time of 100 ms, AGC target of 1e5). The isolation window was 0.7 m/z, and normalized collision energy was 32. Dynamic exclusion was set to 30 seconds and fixed first mass to 100 m/z. Mass spectrometry data were processed using MaxQuant software (version 2.0.1.0). The data were searched against the human reference proteome database (UniProt: UP000005640) with a false discovery rate (FDR) of less than 1% for peptides and proteins. All raw files and MaxQuant output tables have been deposited in the PRIDE repository.

### Data Acquisition and Preprocessing

We note that different datasets use different types of identifiers to refer to a gene or protein. For all datasets we mapped different ID types to HGNC gene symbols and UniProt IDs using the biomaRt and AnnotationDbi R-packages, and updated gene symbols with the HGNChelper package (Durinck et al., 2005; Durinck et al., 2017; Hervé Pagès, 2017; Oh et al., 2022).

#### Dosage Compensation Factor Datasets

We use a series of data sources to identify the factors affecting dosage compensation. In Table 1 we list the datasets, how the data in the dataset has been processed, and their usage for each factor. Missing values of counted factors (e.g., known PTM sites, protein interactions, protein complex participation) were imputed with 0. The factors were combined by an inner join on gene symbol and UniProt ID. Genes with more than 25% of features missing were removed.

#### DNA Copy Number Datasets

For the cell line datasets DepMap and ProCan we obtained absolute gene copy numbers generated with the ABSOLUTE algorithm and chromosome arm copy number alteration (CNA) calls from the DepMap CCLE 23Q2 release (Carter et al., 2012; Cohen-Sharir et al., 2021; DepMap, Broad Institute, 2023). We downloaded the absolute gene copy numbers for the *in vivo* tumor samples from the genomic data commons (GDC) data portal of the National Cancer Institute (NCI) at the National Institutes of Health (NIH), a part of the U.S. Department of Health and Human Services, using the TCGAbiolinks R package (Antonio Colaprico, 2017; Colaprico et al., 2016; Mounir et al., 2019; Silva et al., 2016). The copy numbers were generated using the AscatNGS algorithm which infers the absolute copy number by comparing whole genome sequencing (WGS) data between tumor samples and matched normal samples (Raine et al., 2016). The parameters used for the GDC query were: `project = “CPTAC-3”, workflow.type = “AscatNGS”, data.category = “Copy Number Variation”, data.type = “Gene Level Copy Number”`.

#### Proteomics Datasets

We downloaded pan-cancer cell line proteome and model annotation data from the DepMap CCLE project of the Broad Institute (Dephoure et al., 2014; DepMap, Broad Institute, 2023; DepMap portal; 375 cell lines) and the Cell Model Passports ProCan project of the Wellcome Sanger Institute (Gonçalves et al., 2022; Cell Model Passports portal; 949 cell lines). We downloaded pan-cancer proteome and clinical data for *in vivo* tumor samples from the CPTAC project of the NCI (Li et al., 2023; Proteomic Data Commons; 1026 tumor samples, 523 normal tissue samples; BCM harmonized dataset). We measured the proteome of engineered cell lines in three biological replicates for each RM13, Rtr13, and RPE1p53-/-; this dataset was named P0211 (Figure S7). The dataset of engineered cell lines was created by combining P0211 with previously published data from cell lines: RPE1 p53 KO, RM 10;18, RM 13, RM 19p (Chunduri et al., 2021; PRIDE Accession: PXD018440).

We only used cell line and tumor samples with proteomics data with matched chromosome arm or gene copy number data. From all pan-cancer proteomics data, we removed measurements from proteins encoded on non-autosomal chromosomes. To avoid a disproportional influence of measurement noise, we removed values below 0.1% of protein abundance measurements within each dataset. To ensure the reproducibility of our results across datasets we used the protein-protein reproducibility ranks from Upadhya & Ryan and removed proteins with reproducibility ranks below the 10% percentile (Upadhya & Ryan, 2022). We normalized the measurements between samples within a dataset with cyclic LOESS using the `normalizeBetweenArrays()` function of the `limma` R-package (Ritchie et al., 2015). When analyzing pan-cancer cell lines we only included genes in our buffering analysis that had protein expression data in at least 10 cell lines in which the chromosome arm that encoded this gene was gained, 10 cell lines where this chromosome arm was lost, and 10 cell lines in which this chromosome arm had a neutral ploidy (see Schukken & Sheltzer, 2022). We decreased this threshold to three cell lines per chromosome arm CNA group for the dataset of lab-engineered cell lines.

#### Aneuploidy Score, WGD, CRISPR-KO, Drug Screen, and Mutation Datasets

For cell lines, we obtained aneuploidy scores, whole genome doubling, and ploidy calls from the DepMap portal (Cohen-Sharir et al., 2021; DepMap, Broad Institute, 2023). We estimated the aneuploidy score of tumor samples similar to previous approaches (Bökenkamp et al., 2025; Taylor et al., 2018). Using absolute gene copy numbers obtained for CPTAC, we called gene copy number gain and loss by determining if the copy number was above/below the rounded average ploidy of the tumor sample. We called chromosome arm gain and loss by determining if over 75% of genes encoded on the chromosome arm had copy number gains/losses. We then estimated the aneuploidy score by counting the number of chromosome arm copy number gain and loss events in a tumor sample.

To obtain a list of cancer driver genes we downloaded the cancer gene list from OncoKB^™^ and included genes that were present in at least two sources (https://www.oncokb.org/ cancer-genes; Chakravarty et al., 2017; Suehnholz et al., 2024). Drug sensitivity data and compound metadata was obtained from the PRISM Repurposing screens through the DepMap 23Q2 release (DepMap portal; Corsello et al., 2020; DepMap, Broad Institute, 2023). Gene dependency scores were obtained from gene dependency probability estimates, generated from CRISPR Knock-Out screens using the Chronos algorithm, through the DepMap 23Q2 release (DepMap portal; Dempster et al., 2019, 2021; DepMap, Broad Institute, 2023; Meyers et al., 2017; Pacini et al., 2021). The matrix of damaging mutations obtained through the DepMap 23Q2 release was used to derive TP53 mutation status (DepMap portal; DepMap, Broad Institute, 2023).

### Buffering Ratio

We define the buffering ratio (BR) for quantifying the degree of dosage compensation as

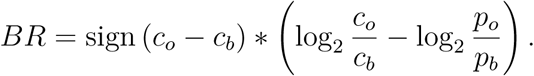

The variables *c_o_* and *p_o_* represent the raw copy number and protein abundance of a gene observed in a sample, while *c_b_ >* 0 and *p_b_ >* 0 are estimated baselines for the copy number and protein abundance of the gene across multiple samples. The sign function obtains the sign (+1 or −1) of the difference *c_o_* − *c_b_*.

#### Confidence Score

The BR is influenced by variability and noise in the proteomics measurement and has by definition an increased sensitivity to changes in the protein abundance, when the change in copy number is small. We defined a confidence score for the BR to estimate the uncertainty of the buffering estimate based on these factors:

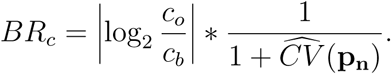

**p_n_** is a vector that contains the protein abundances of a gene in all samples within a dataset where the copy number of that gene was neutral, meaning there was neither a gain or a loss of copy number relative to the copy number baseline. 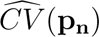 is the sample coefficient of variation (CV) applied to **p_n_**.

### Buffering Classes

#### BR-based

We defined the scaling direction of protein abundance and copy number as positive, if the observed protein abundance and copy number change in the same direction relative to the baseline (formally: (*c_o_* − *c_b_*) ∗ (*p_o_* − *p_b_*) *>* 0), and negative otherwise. A negative scaling direction implies that the protein abundance increased while the copy number decreased relative to the baseline or vice versa. Using protein abundance log2 fold-change (Log2FC) based thresholds, BR based thresholds, and the scaling direction we classified genes as *Scaling*, *Buffered*, or *Anti-Scaling* (Table 2).

**Table 2:**
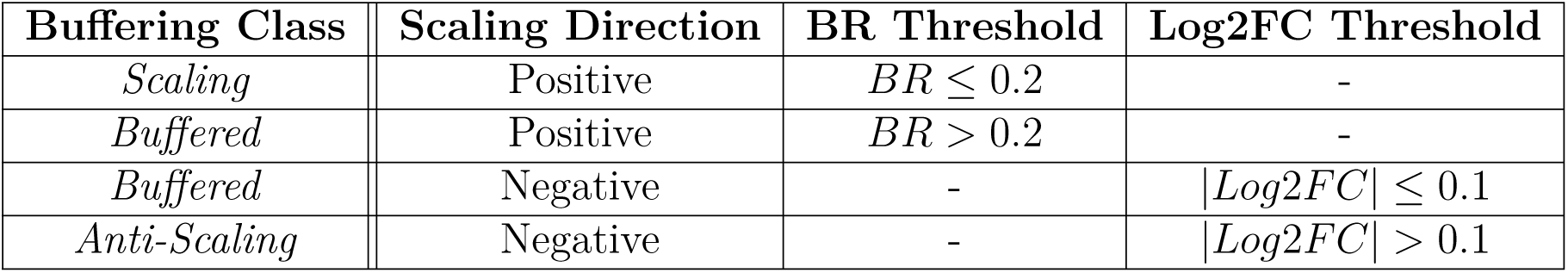
Thresholds used for classifying genes as *Scaling*, *Buffered*, and *Anti-Scaling* using the Buffering Ratio.

#### Log2FC-based

As a control we classified genes as *Scaling*, *Buffered*, or *Anti-Scaling* using Log2FC-based thresholds only as previously published (Table 3; Schukken & Sheltzer, 2022).

**Table 3:**
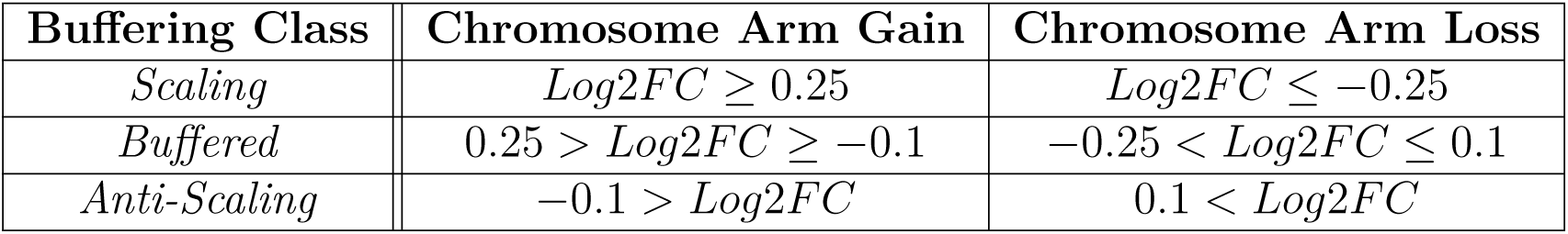
Log2FC thresholds used for classifying genes as *Scaling*, *Buffered*, and *Anti-Scaling*.

| Buffering Class | Chromosome Arm Gain | Chromosome Arm Loss |
| --- | --- | --- |
| *Scaling* | $Log2FC \geq 0.25$ | $Log2FC \leq -0.25$ |
| *Buffered* | $0.25 > Log2FC \geq -0.1$ | $-0.25 < Log2FC \leq 0.1$ |
| *Anti-Scaling* | $-0.1 > Log2FC$ | $0.1 < Log2FC$ |

### Protein Dosage Compensation Analysis

We analyzed protein dosage compensation using both absolute gene copy numbers (CNs, ABSOLUTE algorithm) and chromosome arm copy number alterations (CNAs). We defined four different analysis variants (Table 4) that use different copy number data sources, different classification methods, and produce buffering classes either for each gene in each sample (per-gene, per-sample) or for each gene across all samples (per-gene).

**Table 4:** Variants of the protein dosage compensation analysis.

| Analysis Variant | Granularity | Copy Number Data | Buffering Classes |
| --- | --- | --- | --- |
| GeneCN | per-gene,<br>per-sample | Absolute gene CNs | BR-based |
| ChrArm | per-gene,<br>per-sample | Chromosome arm CNAs | BR-based |
| ChrArm (avg.) | per-gene | Chromosome arm CNAs | Log2FC-based |
| ChrArm (Log2FC) | per-gene,<br>per-sample | Chromosome arm CNAs | Log2FC-based |

For the analysis variant GeneCN, we calculated the copy number and protein abundance baselines *c_b_* and *p_b_* for each gene as the median absolute gene CN and median protein abundance across all samples where the respective chromosome was disomic. We then calculated BR, the confidence score of the BR, and determined BR-based buffering classes using the protein abundances and absolute gene CNs of each gene within each sample. For ChrArm we applied a similar method but used the rounded ploidy of the sample as the copy number baseline *c_b_* and added the chromosome arm CNA (either +1, 0, or −1) to *c_b_* to obtain the observed copy number *c_o_*. ChrArm (Log2FC) uses the same baseline estimation as ChrArm but uses thresholds based on the Log2FC between observed and baseline protein abundance to determine buffering classes. For ChrArm (avg.) we used the method described by Schukken & Sheltzer: For each gene we separated the samples into gain, neutral, and loss cohorts based on the chromosome arm CNA for the chromosome that encodes this specific gene. We then calculated the mean protein abundance Log2FC between gain and neutral, and between loss and neutral cohorts, and applied Log2FC-based thresholds to obtain the buffering classes (Schukken & Sheltzer, 2022).

For the tumor sample dataset CPTAC we applied the GeneCN analysis variant differently by assuming a copy number baseline of *c_b_* = 2 for all genes, using the median abundance of each protein within normal (non-tumor) samples as the protein baseline *p_b_*, using absolute copy numbers generated using the AscatNGS algorithm as observed copy numbers *c_o_*, and using the protein abundance of the corresponding tumor samples as the observed protein abundance *p_o_*. Tumor samples with no corresponding normal samples or tumor samples with a purity below 40% were removed.

### Rank Aggregation

We implemented the mean normalized rank (MNR) to aggregate and unify estimates (e.g. sample BR and ROC AUCs) from different datasets or analysis variants (Upadhya & Ryan, 2022). Let *S*_1_*, . . ., S_n_* be *n* ∈ ℕ sets, each containing *m* features, meaning |*S*_1_| = · · · = |*S_n_*| = *m*. These features can either have present or missing values. For each set *S_i_*, with *i* ∈ {1*, . . ., n*}, we calculated the rank of each feature with value in *S_i_* and divided the rank by the number of features *m*. We then removed features that have more than ⌊*n/*2⌋ missing ranks across all *n* sets. Finally, we obtained the MNR for each feature *f_j_, j* ∈ {1*, . . ., m*} by calculating the arithmetic mean of the ranks of the feature *f_i_* across all sets *S*_1_*, . . ., S_n_*.

### Buffering Ratio Variance Analysis

To determine which genes were consistently buffered, we analyzed the variance of the BR for each gene. For each gene within DepMap, ProCan, and CPTAC, we calculated the mean and standard deviation of the BR (GeneCN analysis variant), the number of samples with an observable BR, the fraction of samples where this gene was classified buffered, and performed a one-sided one-sample Wilcoxon signed-rank test to determine if the BR was significantly above the BR threshold for the *Buffered* class. Within each dataset, we retained genes with more than 20 BR observations, more than 33% of *Buffered* observations, a *Buffered*-fraction higher than the median *Buffered*-fraction of all genes, a mean BR above the *Buffered* threshold (*BR >* 0.2), a significant Benjamini-Hochberg adjusted p-value (*p_adj_ <* 0.05), and a BR standard deviation below 2. We calculated the mean normalized rank (MNR) of the BR standard deviation for each gene across all datasets, and selected 50 genes with the lowest MNR as consistently buffered.

### Sample Buffering Ratio

We defined the sample buffering ratio (sample BR) to quantify the average degree of buffering a sample (aneuploid cell line or tumor sample) exhibits. We removed all non-aneuploid samples, i.e., samples with an aneuploidy score of 0 and a rounded average ploidy of 2. We then removed BR values of genes within a sample with low confidence scores (*BR_c_ <* 0.3) and with a low absolute copy number Log2FC (|log_2_
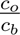| *<* 0.3). Furthermore, we only kept genes that were common across DepMap, ProCan, and CPTAC to ensure comparability of the sample BRs between different proteomics approaches and datasets. Next, we removed all samples that had less than 50 non-missing gene copy number derived BR values left after filtering. Finally, the sample BRs were calculated as the mean gene copy number derived BR (GeneCN analysis variant) across all genes within a sample after filtering. We calculated the z-scores by standardizing the sample BRs within each proteomics dataset they were derived from.

### Analysis of Sample Properties and Cancer Types

Cancer type annotations from DepMap, ProCan, and CPTAC were mapped to OncoTree codes with the same hierarchy (level 2) using the mskcc.oncotree R-package (Kundra et al., 2021). When analyzing sample BR differences between growth patterns (adherent/suspension) we controlled for low aneuploidy by removing cell lines from the adherent cohort with aneuploidy scores above the maximum of the suspension cohort. We controlled for equal aneuploidy by resampling the adherent (target) cohort based on the aneuploidy score distribution in the suspension (reference) cohort. We divided the range of aneuploidy scores into *k* = 6 equal strata by applying the quantile function to the reference cohort aneuploidy scores. We then calculated the fraction of values that fell into each aneuploidy score stratum in the reference and target cohort. For each stratum, we divided the reference sample fraction by the target sample fraction and used this value as the probability for randomly sampling cell lines from the target cohort in the current stratum (without replacement). We then verified that the aneuploidy score difference between the two cohorts is not significantly different after resampling using a Wilcoxon rank-sum (Mann-Whitney U) test. We chose the number of strata to be *k* = 6 using this test condition.

### Unifactorial Analysis

We performed this analysis to understand how well each factor individually predicts whether a gene is buffered or not. We filtered for genes in samples with copy number or chromosome arm gain or loss, by comparing observed copy number against the baseline copy number (for ChrArm, GeneCN) or by filtering for non-zero chromosome arm CNAs (for ChrArm (avg.)). We established a binary classification scheme by removing *Anti-Scaling* observations, setting *Buffered* observations as the case (true) and *Scaling* as the control (false) class. The receiver operating characteristic area under the curve (ROC AUC) was calculated for each factor, within each analysis variant, dataset, and copy number condition (gain, loss) using the pROC package (Robin et al., 2011).

WGD-controlled results for DepMap and ProCan were generated by separating WGD-positive and WGD-negative samples, determining buffering classes for all analysis variants, and calculating unifactorial ROC AUCs as described above. We then calculated mean normalized ranks of the ROC AUCs for each factor and copy number condition (gain, loss) combining all analysis variants across all datasets, all WGD-positive datasets, and all WGD-negative datasets.

#### Bootstrapped Analysis

To determine significant differences of factor ROC AUCs between copy number conditions and analysis variants, we applied filtering and binary classification schemes for each dataset, analysis variant, and copy number condition as described above, bootstrapped the resulting datasets (*n* = 10000 random samples each, with replacement), and calculated ROC AUCs for each factor and bootstrapped dataset. Bootstrapping the ROC AUCs was parallelized using the `parallel` R-package. We calculated the rank correlation of factors between gain and loss conditions within BR-based analysis variants (ChrArm, GeneCN), and between analysis variants within gain and loss conditions using Kendall’s rank correlation coefficient. We removed noisy factors with low predictiveness (median ROC AUC < 0.51) from the rank correlation and further excluded factors that were highly correlated with included factors and contained redundant information (protein/mRNA abundance, activating/repressing transcription factors).

#### BR-Factor Correlation Analysis

Correlations between Buffering Ratio (GeneCN) and factor values for each factor, copy number event, and pan-cancer dataset (DepMap, ProCan, CPTAC) were obtained using Spearman’s rank correlation coefficient and p-values were adjusted by the Benjamini-Yekutieli procedure.

### Multifactorial Analysis

We trained machine learning models to use all available factors for predicting protein buffering. Before training we imputed missing values of each factor with its median value and applied min-max normalization to improve convergence during training. For each dataset, analysis variant, and copy number condition we then filtered for gain or loss, and established a binary classification scheme as described in the unifactorial analysis. The observations in the resulting datasets were then shuffled and split into training and test subsets using a 80/20-split. Models were trained on each of the training datasets using the `caret` R-package with 3-fold cross validation for hyperparameter tuning (Kuhn, 2008). Model performance was evaluated by predicting *Buffered*/*Scaling* labels in the associated test dataset of the model and calculating the ROC AUC on the *Buffered* class probabilities returned by the model. We trained `xgbLinear` (extreme gradient boosting using linear classifiers), `rf` (random forest), and `pcaNNet` (artificial neural network with prior PCA feature transformation) model architectures available in `caret`, however `xgbLinear` produced the highest ROC AUCs in pan-cancer datasets (DepMap, ProCan, CPTAC) among these architectures. Before training we removed factors that were highly correlated, provided redundant information, and confounded the interpretability of the model (e.g. due to overfitting): “Homology Score”, “Protein Abundance”, “mRNA Abundance”, “Transcription Factors (Repressor)”, “Transcription Factors (Activator)”, “Triplosensitivity”.

#### Feature Explanation

We removed observations from a model’s test set where the model’s prediction was incorrect, and calculated SHapley Additive exPlanation values (SHAP values, Lundberg & Lee, 2017) for a random subset of the remaining observations for each feature of the model (300 *Buffered*, 300 *Scaling* observations). To calculate SHAP values, we used a variant of the KernelSHAP algorithm that takes the statistical dependence between features into account and allows for categorically distributed features by using the `ctree` method of the `shapr` R-package (Aas et al., 2021; Redelmeier et al., 2020). We specified *ϕ*_0_, i.e., the expected prediction without any features, as the fraction of *Buffered* observations in the training set of the model, and set the number of sampled feature combinations to 300. Of the trained model architectures, `shapr` only supported calculating SHAP values for `xgbLinear`. To understand what feature values lead to a higher chance of predicting an observation as *Buffered* or *Scaling* when including this feature in the model, we calculated for each feature the Spearman’s correlation coefficient between SHAP values and the associated values of the feature at every observation.

#### Out-of-sample Evaluation

To evaluate the performance of a model across datasets, analysis variants, and copy number conditions, we evaluated the performance of each model on the test set of every other model using the ROC AUC on the *Buffered* class prediction probabilities as described above.

### Differential Expression Analysis

Samples were considered low-buffering if their sample BR was below the 20% percentile of sample BRs and high-buffering if their sample BR was above the 80% percentile. For each gene, we calculated mean protein abundance log_2_ fold-changes (log_2_ *p_high_* − log_2_ *p_low_*), performed unpaired two-tailed Welch’s unequal variances t-tests, and adjusted the p-values using the Benjamini-Hochberg procedure to determine significant protein abundance differences between high- and low-buffering samples. Genes were considered significantly deregulated if |*Log*2*F C*| *>* 0.5 and if the adjusted p-value was below 0.05. We created an additional adherent control dataset by removing all cell lines with non-adherent growth patterns from the ProCan dataset. We then performed splitting and differential expression analysis on this dataset as described above. Genes were called to be commonly deregulated in high/low buffering cell lines if they were significantly up-/downregulated in DepMap, ProCan, and the adherent control.

### Enrichment Analysis

Over-representation analysis was performed using the `gprofiler2` R-package. Plots show the five most enriched terms within each selected term dataset (e.g. KEGG pathway, Gene Ontology, CORUM; Kolberg et al., 2023). We performed gene set enrichment analysis (GSEA) for DepMap, ProCan, and CPTAC using the human hallmark gene sets of the molecular signatures database (MSigDB; Liberzon et al., 2015; Subramanian et al., 2005). Normalized enrichment scores have been generated with the `fGSEA` R-package using the log2 fold-change ranked lists of genes from the differential expression analysis (Korotkevich et al., 2016). We generated aneuploid controls for DepMap, ProCan, and CPTAC by performing differential expression analysis on cell lines separated by aneuploidy score (>80%, <20%) and applying GSEA on the Log2FC ranks. Network analysis of significantly deregulated genes has been performed with STRING using the `STRINGdb` R package (Szklarczyk et al., 2023). We retained edges in the network with a 90% confidence or higher, removed disconnected nodes, and performed clustering using the MCL algorithm with an inflation parameter of 2.

### Proteotoxic Stress Analysis

To estimate the degree of proteotoxic stress per tumor sample, we performed single-sample GSEA (ssGSEA) using the `GSVA` R-package, the MSigDB human hallmark gene sets, and protein abundance measurements from the CPTAC dataset (Hänzelmann et al., 2013). We then calculated Spearman’s rank correlation coefficient between enrichment scores for the `UNFOLDED_PROTEIN_RESPONSE` gene set and the sample BRs of the tumor samples.

### Proliferation Analysis

To estimate the proliferation we obtained the day4-day1 growth ratios from DMSO-treated cell lines in the GDSC drug screen dataset, removed ratios with less than three replicates, and used the ratio with the most replicates for each cell line (Cell Model Passports portal; Iorio et al., 2016). Spearman’s rank correlation coefficient was calculated between day4-to-day1 growth ratios of cell lines, sample BRs calculated for the ProCan and DepMap datasets, and sample BR mean normalized ranks.

### Gene Essentiality Analysis

Differential dependency of buffered cell lines on gene functions was determined using dependency scores generated from CRISPR-KO screens (see Data Acquisition and Pre-processing). We separated cell lines into high- and low-buffering cohorts using 80% and 20% percentile cutoffs on the sample BR mean normalized ranks for cell lines, calculated the mean dependency score difference per gene (*d_high_* − *d_low_*), performed two-tailed Wilcoxon rank-sum (Mann-Whitney U) tests, and adjusted the p-values using the Benjamini-Hochberg procedure. High-/low-buffering cell lines were considered differentially dependent on a gene if the adjusted p-value of the gene was below 0.05 and if |*Log*2*F C*| *>* 0.1. Genes were called exclusively essential in high-buffering cell lines, if the mean dependency score of the gene was significantly higher in high buffering cell lines, its mean dependency score was below 0.5 in low-buffering cell lines, and above 0.5 in high-buffering cell lines. Vice versa, genes were considered exclusively essential in low buffering cell lines for significantly higher dependency scores in low buffering cell lines, a mean dependency score above 0.5 in low buffering cell lines, and below 0.5 in high buffering cell lines.

Similar to the differential expression analysis, we generated an adherent control dataset by re-using the sample BR MNR datset, removing cell lines with non-adherent growth patterns, splitting by sample BR MNR, and performing the differential dependency analysis as described above.

### Drug Sensitivity Analysis

We calculated the sensitivity of high-buffering cells on individual drugs, drug targets and drug mechanisms in two ways. First, we calculated the Spearman correlation coefficient of cell line sample BR mean normalized ranks and median drug effect scores from the PRISM Repurposing screens (see Data Acquisition and Preprocessing) per drug, drug target, and drug mechanism of action. Second, we separated cell lines based on their sample BR MNR (>80%, <20%), calculated the mean drug effect score Log2FC between high- and low-buffering cell lines (log_2_ *e_high_* − log_2_ *e_low_*), and performed unpaired two-tailed Welch’s unequal variances t-tests for each drug, drug target, and drug mechanism. Both correlation and differential drug effect p-values were adjusted using the Benjamini-Hochberg procedure. Mean drug effect score differences were considered significant if the adjusted p-value was below 0.05 and if |*Log*2*F C*| *>* 0.2.

We controlled for aneuploidy by performing differential drug effect analysis on cell lines split by aneuploidy score instead of sample BR MNR percentiles (>80%, <20%). We called drug sensitivities unique for high/low-buffering cell lines, if both correlation and differential drug effect differences were significant and coincided, and if the drug effect differences between high and low aneuploid cell lines were insignificant or significant in the opposite direction.

### Statistical Analysis

Within all boxplots the box depicts the interquartile range with the central band representing the median value of the data. The whiskers represent the furthest datapoint within 1.5 times the interquartile range. Two-tailed Wilcoxon rank-sum (Mann-Whitney U) tests were performed to determine significant differences in BR-derived quantities (e.g. BR, sample BR) when comparing two conditions. Correlations involving BR-derived quantities were calculated using Spearman’s rank correlation coefficient. Unless stated otherwise, tests with *p <* 0.05 were considered significant.

## Acknowledgments

The proteomics mass spectra of the engineered RPE-1 cell lines (P0211) were measured by the Center for MS Analytics at RPTU Kaiserslautern (https://bio.rptu.de/center-ms-analytics). This project was supported by grant projects BioComp 4.0 and MLKL (funded by Research Initiative of Rhineland-Palatine to Z.S.). Research in the Sheltzer Lab is supported by multiple sources, including NIH grants R01CA237652 and R01CA276666, an American Cancer Society Research Scholar Grant, a grant from Project 8p, grant #2023236 from the US-Israel Binational Science Foundation, a grant from the Andrew McDonough B+ Foundation, and a sponsored research agreement from Ono Pharmaceutical.

## Author contributions

Z.S., J.M.S., K.M.S., and E.M.H. conceived the idea, E.M.H. performed the study, K.B. and M.R. performed the proteomics experiment, E.M.H. and Z.S. wrote the manuscript, all authors read and commented on the manuscript.

## Disclosure and competing interest statement

J.M.S. is an inventor on a patent related to chromosome engineering and the construction of aneuploid genomes. J.M.S. is a co-founder of and equity holder in Meliora Therapeutics and KaryoVerse Therapeutics.

## Data Availability

The datasets and computer code produced in this study are available in the following databases:

- Mass spectrometry proteomics data of the chromosome engineered RPE-1 cell lines (P0211): PRIDE PXD060017 (https://www.ebi.ac.uk/pride/archive/projects/PXD060017, username, password: NypjXh6UGyYq)
- Computational study code: GitHub (https://github.com/sheltzer-lab/DosageCompensationFactors)

## Supplemental Figures

**Figure S1:**
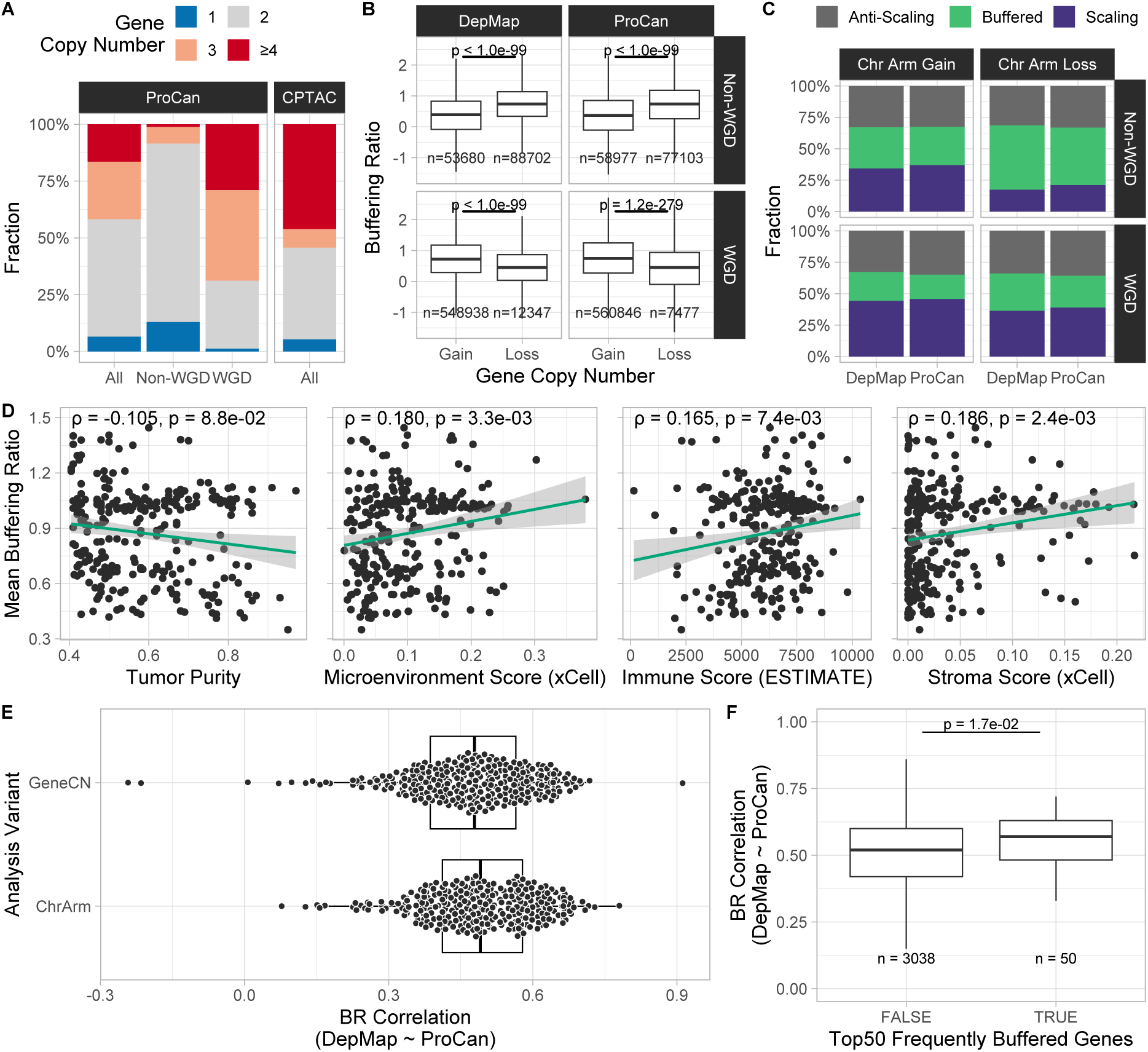
Controlling for whole genome doubling status and tumor purity confirms observed protein buffering patterns. (A) Distribution of gene copy numbers among cell lines (ProCan) and tumor samples (CPTAC) separated by their whole genome doubling (WGD) status. (B) Difference in buffering ratio distribution of all proteins between gene copy number gain and loss sep-arated by the WGD status. (C) Categorical distribution of buffering classes across pan-cancer cell line datasets upon gene copy number gain and loss separated by WGD status. (D) Correlation between mean buffering ratio per tumor sample (CPTAC) and purity scores obtained from metadata provided by CPTAC (Spearman’s *ρ*). (E) Correlation of buffering ratios per cell line between DepMap and ProCan (Spearman’s *ρ*), separated by buffering ratios created using either gene (GeneCN) or chromosome arm copy numbers (ChrArm). (F) Gene-wise buffering ratio correlation between DepMap and ProCan (GeneCN, Spearman’s *ρ*), separated by their presence in the list of frequently buffered genes with low BR variance.

**Figure S2:**
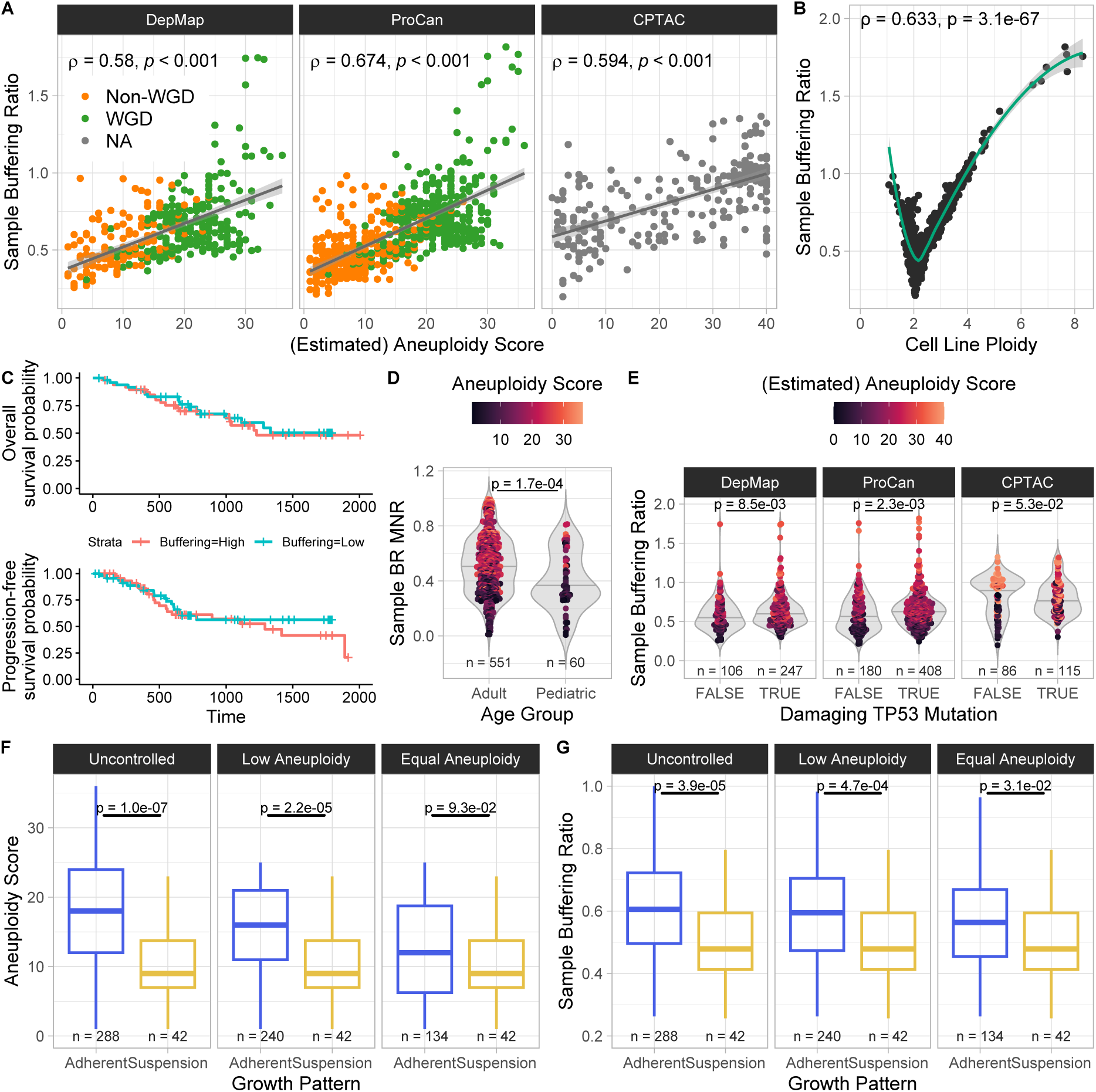
Average buffering per sample shows relationship with cellular and clinical features of cancer. **(A)** Scatter plot showing a positive correlation between sample buffering ratio and aneuploidy score in all pan-cancer datasets (Spearman’s *ρ*). **(B)** Scatter plot showing a LOESS-smoothed non-linear relationship between sample buffering ratio and cell line ploidy (ProCan). **(C)** Kaplan-Meyer plots showing a decreased progression-free and overall survival probability for tumor samples with high buffering (>80% quantile of sample buffering ratio) compared to low buffering (<20% quantile, CPTAC). **(D)** Difference in sample buffering ratio mean normalized ranks between age categories shows decreased sample-wide buffering in pediatric cancers (Wilcoxon rank-sum test). **(E)** Difference in sample buffering ratio between samples with and without damaging TP53 mutations (Wilcoxon rank-sum test). **(F)** Difference in aneuploidy between cell lines grown in suspension and adherent cell culture types when applying aneuploidy control methods (DepMap). Controlled for aneuploidy by removing cell lines from the adherent cohort with aneuploidy scores above the maximum of the suspension cohort (middle), and by using stratified sampling to ensure an equal distribution of aneuploidy scores (right). **(G)** Difference in sample buffering ratio between cell lines grown in suspension and adherent cell culture types, controlled for aneuploidy (DepMap).

**Figure S3:**
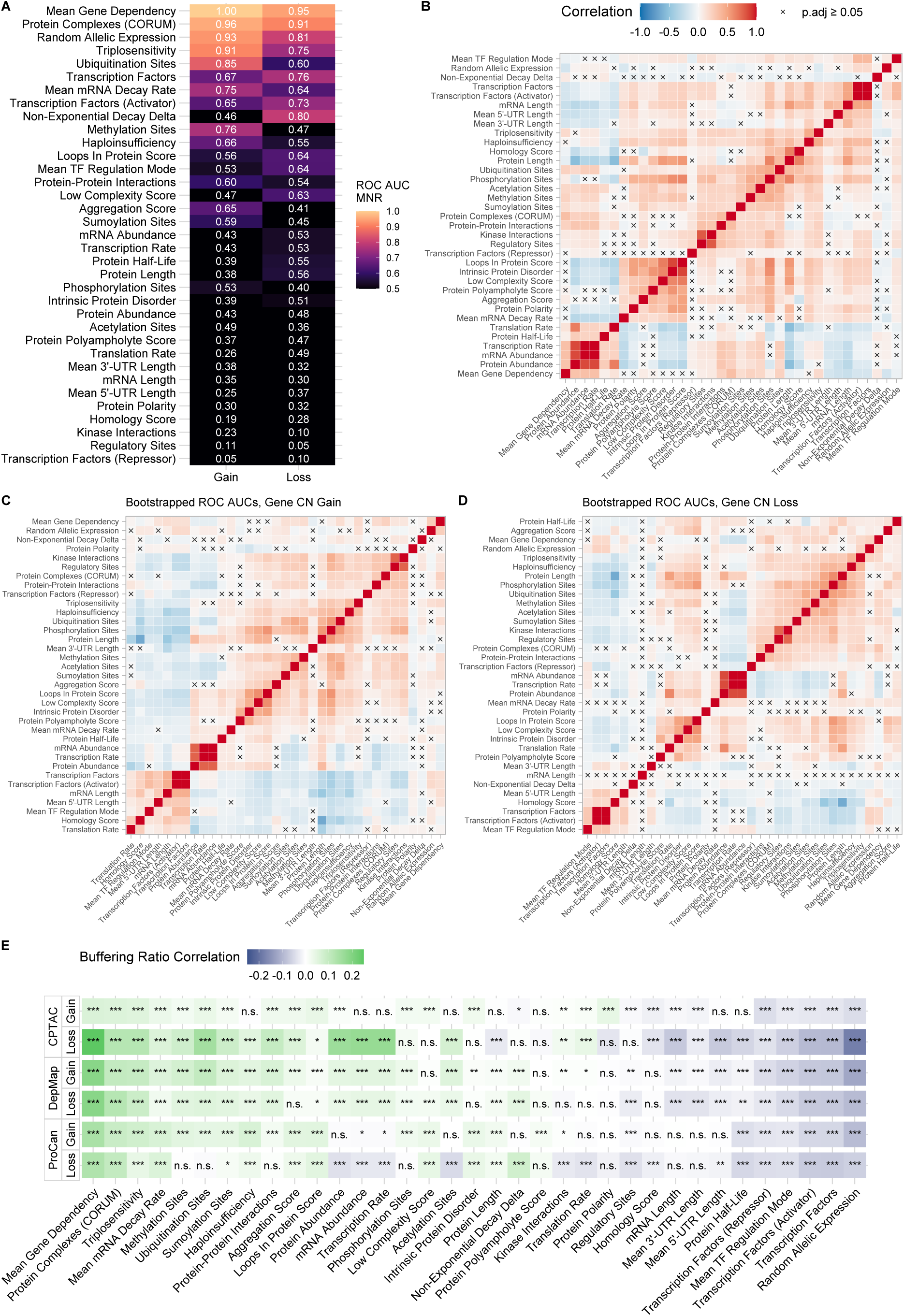
Correlation patterns of potential dosage compensation factors. **(A)** Mean normalized ranks (MNRs) of ROC AUCs of each factor used for classifying whether a gene is *Buffered* on protein level or *Scaling*, while excluding the *Anti-Scaling* class. MNRs were calculated across all analysis variants (GeneCN, ChrArm, ChrArmAvg) and pan-cancer datasets (DepMap, ProCan, CPTAC), grouped by gene or chromosome arm copy number gain and loss. WGD-controlled subsets were excluded. **(B-D)** Clustered heatmaps showing the correlation (Spearman’s *ρ*) between factors. Correlations with insignificant Benjamini-Hochberg adjusted p-values were crossed out (*p_adj_* ≥ 0.05). Correlations were calculated using factor values **(B)**, bootstrapped ROC AUCs for predicting protein buffering upon gene copy number gain **(C)**, and bootstrapped ROC AUCs upon gene copy number loss **(D)**. **(D)** Spearman correlation between factors values and buffering ratios upon gene copy number loss and gain across all pan-cancer datasets (DepMap, ProCan, CPTAC).

**Figure S4:**
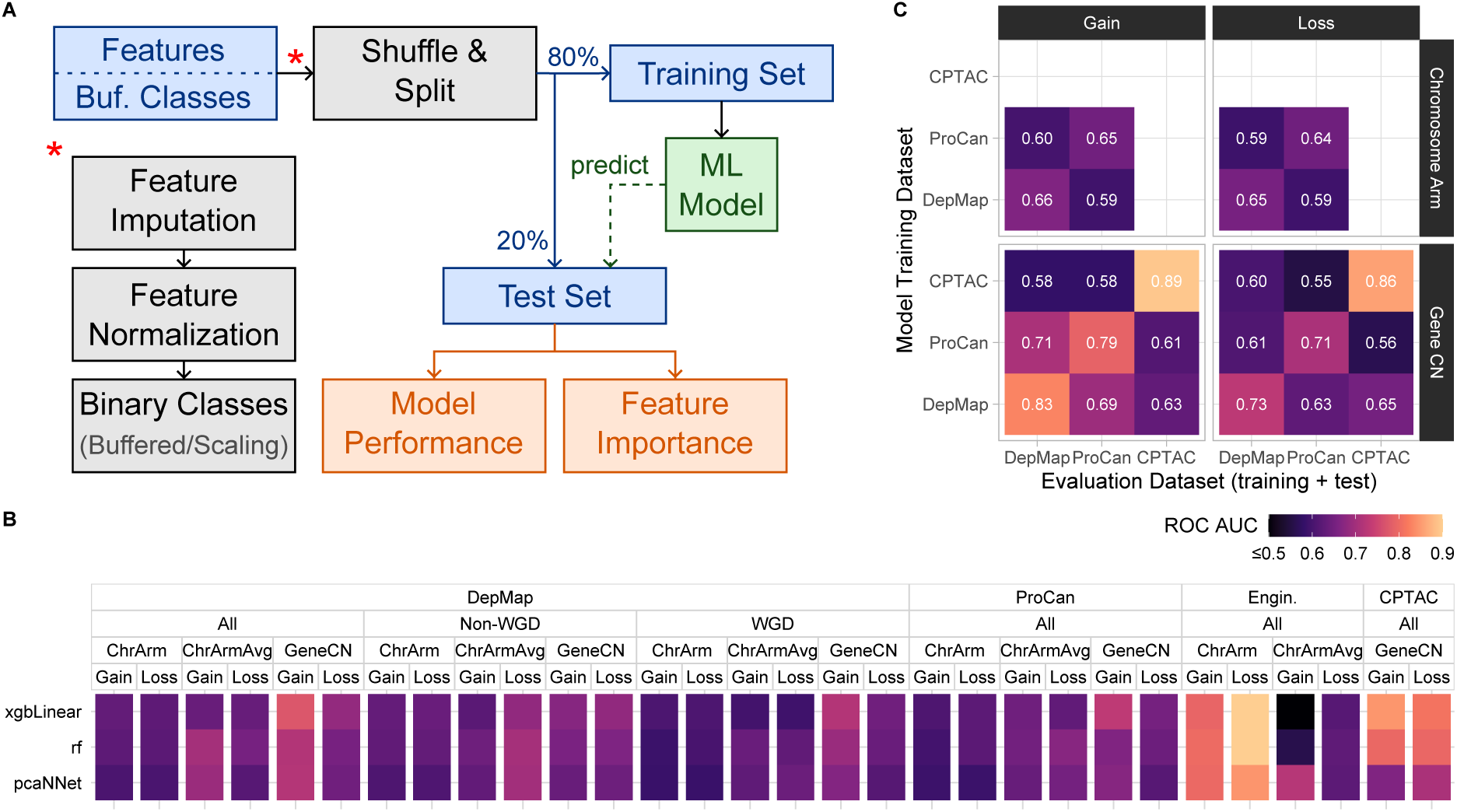
Multifactorial models improve prediction performance of protein buffering. **(A)** Illustration of training and evaluation procedure of multifactorial models. **(B)** Model performance (ROC AUC) of trained models, separated by model architecture (xgbLinear, rf, pcaNNet), dataset (DepMap, ProCan, CPTAC, Engineered), subset (All, WGD, Non-WGD), analysis variant (GeneCN, ChrArm, ChrArmAvg), and copy number event (Gain, Loss). **(C)** Out-of-sample performance of multifactorial models trained on buffering classes derived from gene and chromosome arm copy number gain and loss data from different pan-cancer datasets (DepMap, ProCan, CPTAC). Model performance was evaluated using the ROC AUC of model predictions on merged training and test datasets.

**Figure S5:**
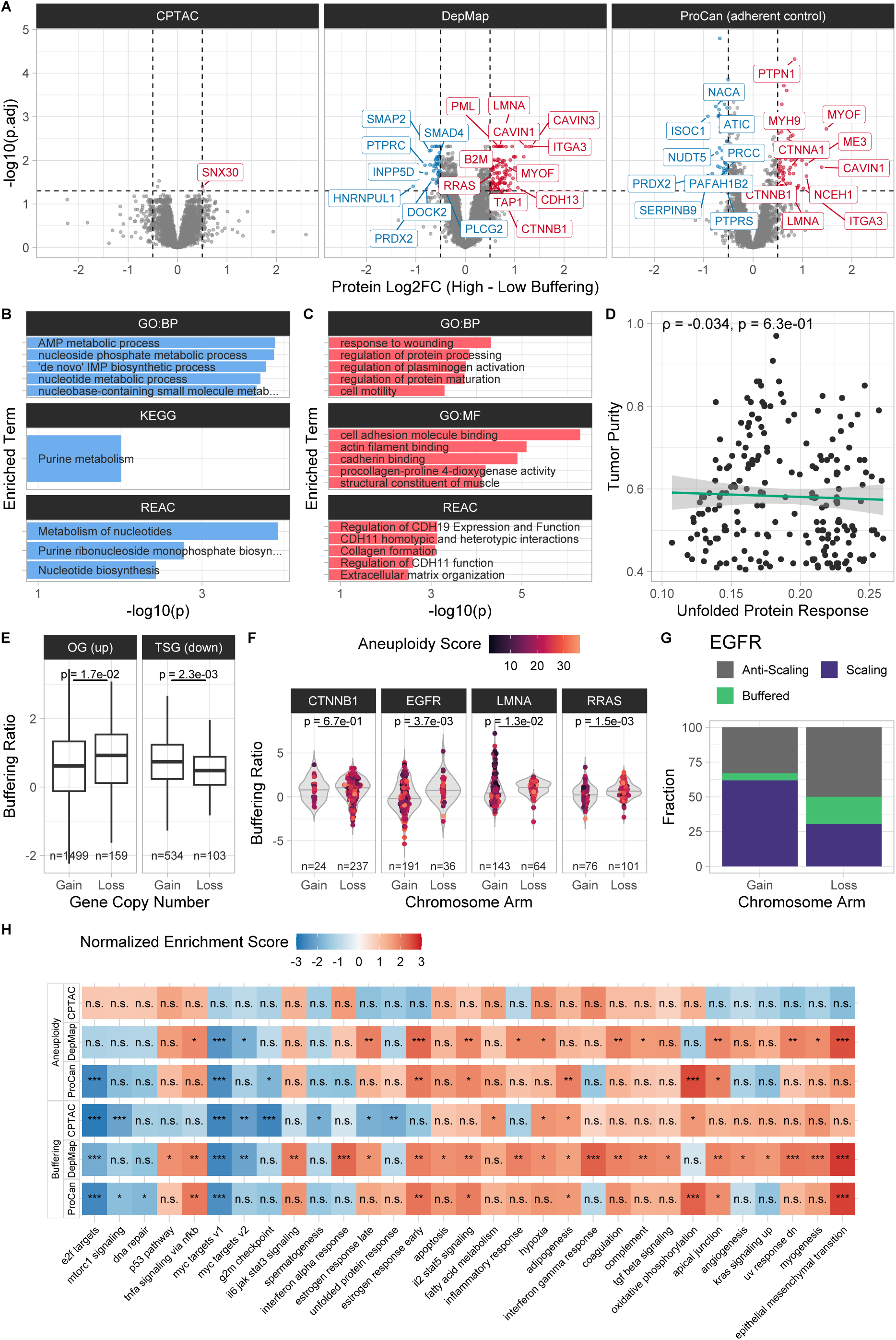
Enrichment patterns suggest oncogenic effect of protein buffering. **(A)** Volcano plots showing log2 fold-change between high (>80% quantile of sample buffering ratios) and low (<20% quantile) buffering cell lines and tumor samples in DepMap, CPTAC, and adherent control datasets (Student’s t-test, Benjamini-Hochberg adjusted p-values, significance thresholds: |*Log*2*F C*| *>* 0.5, *p_adj_ <* 0.05). **(B, C)** Over-representation analysis (ORA) of genes that are significantly down-**(B)** and up-regulated **(C)** in highly buffering cell lines with adherent growth patterns (ProCan, adherent con-trol). **(D)** Correlation between tumor purity and single-sample enrichment scores for the *Unfolded Protein Response* HALLMARK gene set using CPTAC tumor sample pro-teome data. **(E)** Buffering ratio distribution of oncogenes (OG) significantly upregulated in high buffering cell lines and of downregulated tumor supressor genes (TSG) upon gene copy number gain and loss (ProCan, all cell lines). **(F)** Buffering ratio distribu-tion of selected oncogenes upon gene copy number gain and loss (ProCan). **(G)** Cat-egorical distribution of buffering classes for EGFR upon gain and loss of chromosome arm 7p (ProCan). **(H)** Normalized enrichment scores of MSigDB HALLMARK gene sets for pan-cancer datasets (DepMap, ProCan, CPTAC). Enrichment scores have been generated by comparing high against low buffering samples and comparing high against low aneuploid samples (>80% and <20% quantiles of sample buffering ratio and an-euploidy score estimates). Significance levels of Benjamini-Hochberg adjusted p-values: (∗)*p_adj_ <* 0.01, (∗∗)*p_adj_ <* 0.001, (∗ ∗ ∗)*p_adj_ <* 0.0001. Removed gene sets with no signi-ficant enrichment scores.

**Figure S6:**
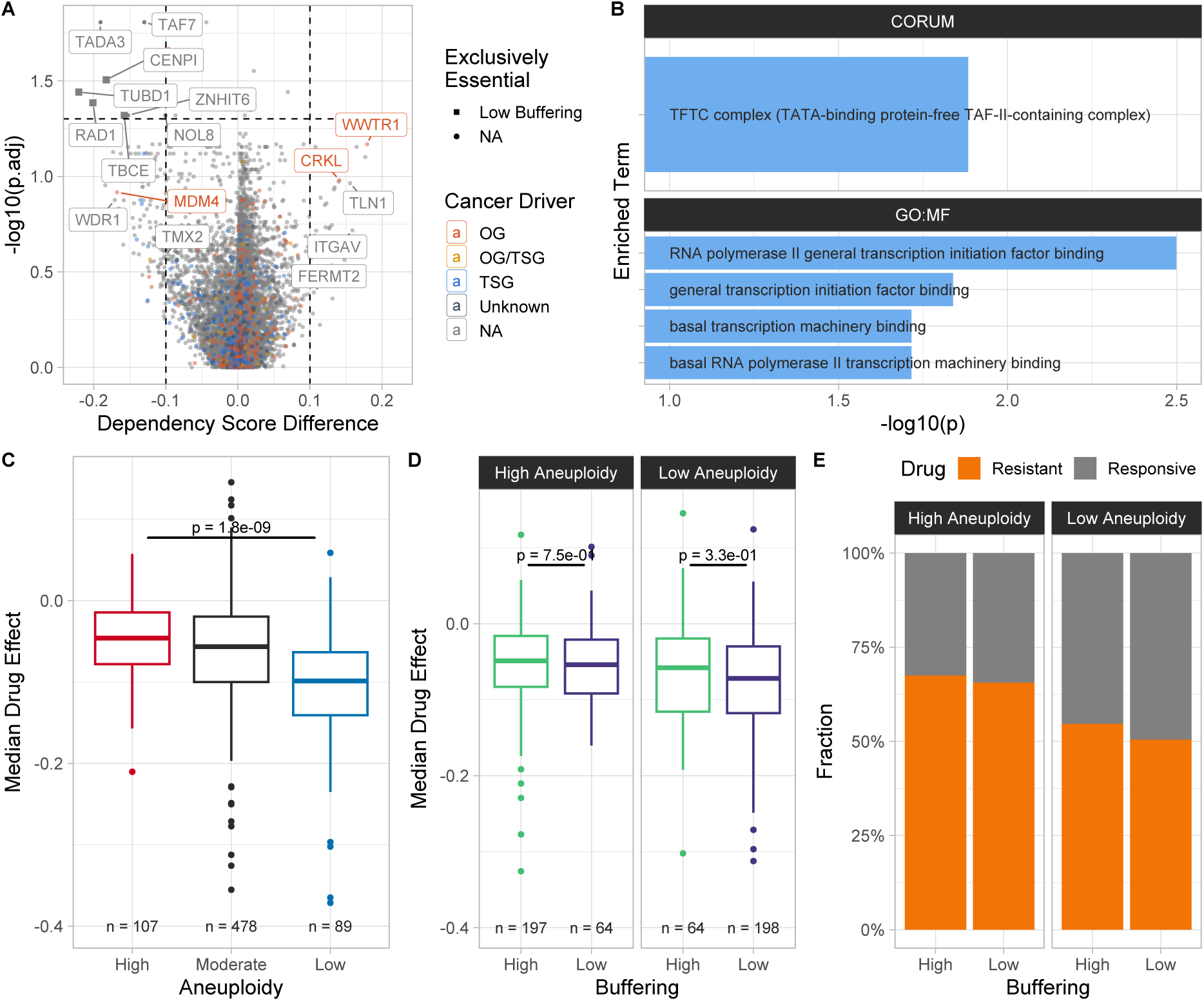
Adherent and aneuploid controls for CRISPR knock-out and drug sensitivity analyses. **(A)** Volcano plot showing the difference of the CRISPR-KO dependency score between high (>80% quantile of sample BR MNRs) and low (<20% quantile) buffering cell lines plotted against Benjamini-Hochberg adjusted Wilcoxon rank-sum test p-values. Cell lines without adherent growth patterns were removed prior to the analysis. Genes were highlighted if they were significant in the adherent control or among the top10 significant genes in Figure 6A. **(B)** Over-representation analysis (ORA) of genes with significantly reduced CRISPR-KO dependency scores in highly buffering cell lines with adherent growth patterns. **(C)** Median drug effect on cell viability across drugs from the PRISM Repurposing dataset for cell lines high (>80% quantile of aneuploidy score), moderate (20%-80%), and low aneuploidy (<20%). **(D)** Median drug effect for high (>50% sample BR MNR) and low buffering cell lines, separated by median aneuploidy score. **(E)** Categorical distribution of drug resistant (>50% median drug effect) and responsive cells between high (>50% sample BR MNR) and low buffering cell lines, separated by median aneuploidy score.

**Figure S7:**
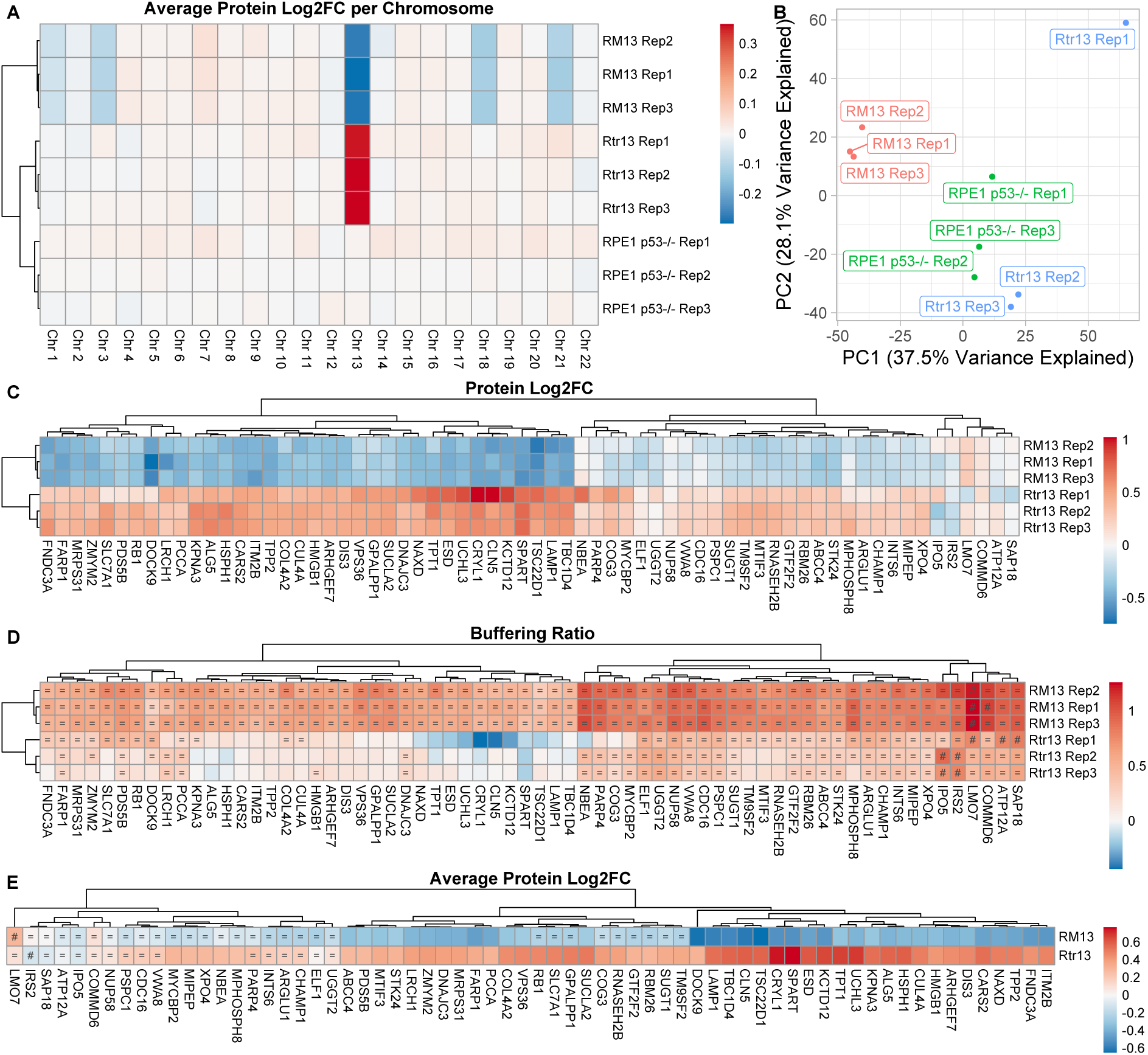
Proteome and protein buffering analysis of RPE-1 cell lines with engineered aneuploidies on chromosome 13 (P0211). **(A)** Mean log2 fold-change of protein abundance per chromosome between cell line rep-licates and disomic baseline (median protein abundance of unaltered RPE-1 replicates per protein). **(B)** Principal components analysis (PCA) of P0211 replicates after normaliz-ation (first two principal components are displayed). **(C)** Protein abundance log2 fold-change of proteins encoded on chromosome 13 between aneuploid replicates and disomic baseline. **(D)** Chromosome arm copy number-derived buffering ratios (BR) of proteins encoded on chromosome 13 in aneuploid replicates. Proteins are classified as *Buffered* (=) and *Anti-Scaling* (#) using BR-based thresholds (see Methods). **(E)** Mean protein abundance log2 fold-changes of proteins encoded on chromosome 13 between aneuploid and non-aneuploid cell lines. Proteins were classified as *Buffered* (=) and *Anti-Scaling* (#) using Log2FC-based thresholds (see Methods).

## References

Aas, K., Jullum, M., & Løland, A. (2021). Explaining individual predictions when features are dependent: More accurate approximations to shapley values. Artificial Intelligence, 298, 103502. 10.1016/j.artint.2021.103502

Adams, B. M., Canniff, N. P., Guay, K. P., Larsen, I. S. B., & Hebert, D. N. (2020). Quantitative glycoproteomics reveals cellular substrate selectivity of the ER protein quality control sensors UGGT1 and UGGT2. eLife, 9, e63997. 10.7554/eLife.63997

Alanis-Lobato, G., Andrade-Navarro, M. A., & Schaefer, M. H. (2017). HIPPIE v2.0: Enhancing meaningfulness and reliability of protein–protein interaction networks. Nucleic Acids Res, 45, D408–D414. 10.1093/nar/gkw985

Andrade, J. R., Gallagher, A. D., Maharaj, J., & McClelland, S. E. (2023). Disentangling the roles of aneuploidy, chromosomal instability and tumour heterogeneity in developing resistance to cancer therapies. Chromosome Res, 31(4), 28. 10.1007/s10577-023-09737-5

Antonio Colaprico, T. C. S. (2017). TCGAbiolinks. Bioconductor. 10.18129/B9.BIOC.TCGABIOLINKS

Badia-i-Mompel, P., Vélez Santiago, J., Braunger, J., Geiss, C., Dimitrov, D., Müller-Dott, S., Taus, P., Dugourd, A., Holland, C. H., Ramirez Flores, R. O., & Saez-Rodriguez, J. (2022). decoupleR: Ensemble of computational methods to infer biological activities from omics data (M. L. Kuijjer, Ed.). Bioinformatics Advances, 2(1), vbac016. 10.1093/bioadv/vbac016

Barbie, D. A., Tamayo, P., Boehm, J. S., Kim, S. Y., Moody, S. E., Dunn, I. F., Schinzel, A. C., Sandy, P., Meylan, E., Scholl, C., Fröhling, S., Chan, E. M., Sos, M. L., Michel, K., Mermel, C., Silver, S. J., Weir, B. A., Reiling, J. H., Sheng, Q., . . . Hahn, W. C. (2009). Systematic RNA interference reveals that oncogenic KRAS-driven cancers require TBK1. Nature, 462(7269), 108–112. 10.1038/nature08460

Bochman, M. L., & Schwacha, A. (2009). The mcm complex: Unwinding the mechanism of a replicative helicase. Microbiol Mol Biol Rev, 73(4), 652–683. 10.1128/MMBR.00019-09

Bökenkamp, J.-E., Keuper, K., Redel, S., Barthel, K., Johnson, L., Becker, A., Wieland, A., Räschle, M., & Storchová, Z. (2025). Proteogenomic analysis reveals adaptive strategies for alleviating the consequences of aneuploidy in cancer. EMBO J, 44(6), 1829–1865. 10.1038/s44318-025-00372-w

Brembeck, F. H., Rosário, M., & Birchmeier, W. (2006). Balancing cell adhesion and wnt signaling, the key role of β-catenin. Current Opinion in Genetics & Development, 16(1), 51–59. 10.1016/j.gde.2005.12.007

Brennan, C. M., Vaites, L. P., Wells, J. N., Santaguida, S., Paulo, J. A., Storchova, Z., Harper, J. W., Marsh, J. A., & Amon, A. (2019). Protein aggregation mediates stoichiometry of protein complexes in aneuploid cells. Genes Dev., 33(15), 1031– 1047. 10.1101/gad.327494.119

Carter, S. L., Cibulskis, K., Helman, E., McKenna, A., Shen, H., Zack, T., Laird, P. W., Onofrio, R. C., Winckler, W., Weir, B. A., Beroukhim, R., Pellman, D., Levine, D. A., Lander, E. S., Meyerson, M., & Getz, G. (2012). Absolute quantification of somatic DNA alterations in human cancer. Nat Biotechnol, 30(5), 413–421. 10.1038/nbt.2203

Chakravarty, D., Gao, J., Phillips, S., Kundra, R., Zhang, H., Wang, J., Rudolph, J. E., Yaeger, R., Soumerai, T., Nissan, M. H., Chang, M. T., Chandarlapaty, S., Traina, T. A., Paik, P. K., Ho, A. L., Hantash, F. M., Grupe, A., Baxi, S. S., Callahan, M. K., . . . Schultz, N. (2017). OncoKB: A precision oncology knowledge base. JCO Precision Oncology, (1), 1–16. 10.1200/PO.17.00011

Chunduri, N. K., Menges, P., Zhang, X., Wieland, A., Gotsmann, V. L., Mardin, B. R., Buccitelli, C., Korbel, J. O., Willmund, F., Kschischo, M., Raeschle, M., & Storchova, Z. (2021). Systems approaches identify the consequences of monosomy in somatic human cells. Nat Commun, 12(1), 5576. 10.1038/s41467-021-25288-x

Ciryam, P., Tartaglia, G. G., Morimoto, R. I., Dobson, C. M., & Vendruscolo, M. (2013). Widespread aggregation and neurodegenerative diseases are associated with super-saturated proteins. Cell Reports, 5(3), 781–790. 10.1016/j.celrep.2013.09.043

Cohen-Sharir, Y., McFarland, J. M., Abdusamad, M., Marquis, C., Bernhard, S. V., Kazachkova, M., Tang, H., Ippolito, M. R., Laue, K., Zerbib, J., Malaby, H. L. H., Jones, A., Stautmeister, L.-M., Bockaj, I., Wardenaar, R., Lyons, N., Nagaraja, A., Bass, A. J., Spierings, D. C. J., . . . Ben-David, U. (2021). Aneuploidy renders cancer cells vulnerable to mitotic checkpoint inhibition. Nature, 590(7846), 486–491. 10.1038/s41586-020-03114-6

Colaprico, A., Silva, T. C., Olsen, C., Garofano, L., Cava, C., Garolini, D., Sabedot, T. S., Malta, T. M., Pagnotta, S. M., Castiglioni, I., Ceccarelli, M., Bontempi, G., & Noushmehr, H. (2016). TCGAbiolinks: An r/bioconductor package for integrative analysis of TCGA data. Nucleic Acids Research, 44(8), e71–e71. 10.1093/nar/gkv1507

Collins, R. L., Glessner, J. T., Porcu, E., Lepamets, M., Brandon, R., Lauricella, C., Han, L., Morley, T., Niestroj, L.-M., Ulirsch, J., Everett, S., Howrigan, D. P., Boone, P. M., Fu, J., Karczewski, K. J., Kellaris, G., Lowther, C., Lucente, D., Mohajeri, K., . . . Esko, T. (2022). A cross-disorder dosage sensitivity map of the human genome. Cell, 185(16), 3041–3055.e25. 10.1016/j.cell.2022.06.036

Corsello, S. M., Nagari, R. T., Spangler, R. D., Rossen, J., Kocak, M., Bryan, J. G., Humeidi, R., Peck, D., Wu, X., Tang, A. A., Wang, V. M., Bender, S. A., Lemire, E., Narayan, R., Montgomery, P., Ben-David, U., Garvie, C. W., Chen, Y., Rees, M. G., . . . Golub, T. R. (2020). Discovering the anticancer potential of non-oncology drugs by systematic viability profiling. Nat Cancer, 1(2), 235–248. 10.1038/s43018-019-0018-6

Davoli, T., Xu, A. W., Mengwasser, K. E., Sack, L. M., Yoon, J. C., Park, P. J., & Elledge, S. J. (2013). Cumulative haploinsufficiency and triplosensitivity drive aneuploidy patterns and shape the cancer genome. Cell, 155(4), 948–962. 10.1016/j.cell.2013.10.011

Dempster, J. M., Boyle, I., Vazquez, F., Root, D. E., Boehm, J. S., Hahn, W. C., Tsherniak, A., & McFarland, J. M. (2021). Chronos: A cell population dynamics model of CRISPR experiments that improves inference of gene fitness effects. Genome Biol, 22(1), 343. 10.1186/s13059-021-02540-7

Dempster, J. M., Rossen, J., Kazachkova, M., Pan, J., Kugener, G., Root, D. E., & Tsherniak, A. (2019, July 31). Extracting biological insights from the project achilles genome-scale CRISPR screens in cancer cell lines [Preprint, bioRxiv]. 10.1101/720243

Dephoure, N., Hwang, S., O’Sullivan, C., Dodgson, S. E., Gygi, S. P., Amon, A., & Torres, E. M. (2014). Quantitative proteomic analysis reveals posttranslational responses to aneuploidy in yeast. eLife, 3, e03023. 10.7554/eLife.03023

DepMap, Broad Institute. (2023). DepMap 23q2 public [DATASET]. 10.6084/M9.FIGSHARE.22765112.V4

Donnelly, N., Passerini, V., Dürrbaum, M., Stingele, S., & Storchová, Z. (2014). HSF1 deficiency and impaired HSP90-dependent protein folding are hallmarks of aneuploid human cells. EMBO J, 33(20), 2374–2387. 10.15252/embj.201488648

Donnelly, N., & Storchová, Z. (2015). Aneuploidy and proteotoxic stress in cancer. Molecular & Cellular Oncology, 2(2), e976491. 10.4161/23723556.2014.976491

Durinck, S., Moreau, Y., Kasprzyk, A., Davis, S., De Moor, B., Brazma, A., & Huber, W. (2005). BioMart and bioconductor: A powerful link between biological databases and microarray data analysis. Bioinformatics, 21(16), 3439–3440. 10.1093/bioinformatics/bti525

Durinck, S., Huber, W., Davis, S., & Smith, M. (2017). biomaRt. Bioconductor. 10.18129/B9.BIOC.BIOMART

Fukushi, J.-i., Makagiansar, I. T., & Stallcup, W. B. (2004). NG2 proteoglycan promotes endothelial cell motility and angiogenesis via engagement of galectin-3 and α3β1 integrin. MBoC, 15(8), 3580–3590. 10.1091/mbc.e04-03-0236

Garcia-Alonso, L., Holland, C. H., Ibrahim, M. M., Turei, D., & Saez-Rodriguez, J. (2019). Benchmark and integration of resources for the estimation of human transcription factor activities. Genome Res., 29(8), 1363–1375. 10.1101/gr.40663.118

Gestaut, D., Limatola, A., Joachimiak, L., & Frydman, J. (2019). The ATP-powered gymnastics of TRiC/CCT: An asymmetric protein folding machine with a symmetric origin story. Current Opinion in Structural Biology, 55, 50–58. 10.1016/j.sbi.2019.03.002

Ghandi, M., Huang, F. W., Jané-Valbuena, J., Kryukov, G. V., Lo, C. C., McDonald, E. R., Barretina, J., Gelfand, E. T., Bielski, C. M., Li, H., Hu, K., Andreev-Drakhlin, A. Y., Kim, J., Hess, J. M., Haas, B. J., Aguet, F., Weir, B. A., Rothberg, M. V., Paolella, B. R., . . . Sellers, W. R. (2019). Next-generation characterization of the cancer cell line encyclopedia. Nature, 569(7757), 503–508. 10.1038/s41586-019-1186-3

Ghosh, P., Beas, A. O., Bornheimer, S. J., Garcia-Marcos, M., Forry, E. P., Johannson, C., Ear, J., Jung, B. H., Cabrera, B., Carethers, J. M., & Farquhar, M. G. (2010). A gαi–GIV molecular complex binds epidermal growth factor receptor and determines whether cells migrate or proliferate (J. S. Gutkind, Ed.). MBoC, 21(13), 2338–2354. 10.1091/mbc.e10-01-0028

Giurgiu, M., Reinhard, J., Brauner, B., Dunger-Kaltenbach, I., Fobo, G., Frishman, G., Montrone, C., & Ruepp, A. (2019). CORUM: The comprehensive resource of mammalian protein complexes—2019. Nucleic Acids Research, 47, D559–D563. 10.1093/nar/gky973

Gonçalves, E., Fragoulis, A., Garcia-Alonso, L., Cramer, T., Saez-Rodriguez, J., & Beltrao, P. (2017). Widespread post-transcriptional attenuation of genomic copy-number variation in cancer. Cell Systems, 5(4), 386–398.e4. 10.1016/j.cels.2017.08.013

Gonçalves, E., Poulos, R. C., Cai, Z., Barthorpe, S., Manda, S. S., Lucas, N., Beck, A., Bucio-Noble, D., Dausmann, M., Hall, C., Hecker, M., Koh, J., Lightfoot, H., Mahboob, S., Mali, I., Morris, J., Richardson, L., Seneviratne, A. J., Shepherd, R., . . . Reddel, R. R. (2022). Pan-cancer proteomic map of 949 human cell lines. Cancer Cell, 40(8), 835–849.e8. 10.1016/j.ccell.2022.06.010

Gwon, Y., Maxwell, B. A., Kolaitis, R.-M., Zhang, P., Kim, H. J., & Taylor, J. P. (2021). Ubiquitination of g3bp1 mediates stress granule disassembly in a context-specific manner. Science, 372(6549), eabf6548. 10.1126/science.abf6548

Hänzelmann, S., Castelo, R., & Guinney, J. (2013). GSVA: Gene set variation analysis for microarray and RNA-seq data. BMC Bioinformatics, 14(1), 7. 10.1186/1471-2105-14-7

Hausser, J. (2019, August 3). Central dogma rates and the trade-off between precision and economy [DATASET]. 10.17632/2VBRG3W4P3.1

Hausser, J., Mayo, A., Keren, L., & Alon, U. (2019). Central dogma rates and the trade-off between precision and economy in gene expression. Nat Commun, 10(1), 68. 10.1038/s41467-018-07391-8

Hervé Pagès, M. C. (2017). AnnotationDbi. Bioconductor. 10.18129/B9.BIOC.ANNOTATIONDBI

Hoege, C., Pfander, B., Moldovan, G.-L., Pyrowolakis, G., & Jentsch, S. (2002). RAD6-dependent DNA repair is linked to modification of PCNA by ubiquitin and SUMO. Nature, 419(6903), 135–141. 10.1038/nature00991

Hornbeck, P. V., Zhang, B., Murray, B., Kornhauser, J. M., Latham, V., & Skrzypek, E. (2015). PhosphoSitePlus, 2014: Mutations, PTMs and recalibrations. Nucleic Acids Research, 43, D512–D520. 10.1093/nar/gku1267

Hwang, S., Cavaliere, P., Li, R., Zhu, L. J., Dephoure, N., & Torres, E. M. (2021). Consequences of aneuploidy in human fibroblasts with trisomy 21. Proc. Natl. Acad. Sci. U.S.A., 118(6), e2014723118. 10.1073/pnas.2014723118

Iorio, F., Knijnenburg, T. A., Vis, D. J., Bignell, G. R., Menden, M. P., Schubert, M., Aben, N., Gonçalves, E., Barthorpe, S., Lightfoot, H., Cokelaer, T., Greninger, P., van Dyk, E., Chang, H., de Silva, H., Heyn, H., Deng, X., Egan, R. K., Liu, Q., . . . Garnett, M. J. (2016). A landscape of pharmacogenomic interactions in cancer. Cell, 166(3), 740–754. 10.1016/j.cell.2016.06.017

Ippolito, M. R., Zerbib, J., Eliezer, Y., Reuveni, E., Vigano, S., De Feudis, G., Shulman, E. D., Savir Kadmon, A., Slutsky, R., Chang, T., Campagnolo, E. M., Taglietti, S., Scorzoni, S., Gianotti, S., Martin, S., Muenzner, J., Mulleder, M., Rozenblum, N., Rubolino, C., . . . Ben-David, U. (2024). Increased RNA and protein degradation is required for counteracting transcriptional burden and proteotoxic stress in human aneuploid cells. Cancer Discovery, 14(12), 2532–2553. 10.1158/2159-8290.CD-23-0309

Ishikawa, K., Makanae, K., Iwasaki, S., Ingolia, N. T., & Moriya, H. (2017). Post-translational dosage compensation buffers genetic perturbations to stoichiometry of protein complexes (A. K. Hopper, Ed.). PLoS Genet, 13(1), e1006554. 10.1371/journal.pgen.1006554

Jensen, S. S., Aaberg-Jessen, C., Christensen, K. G., & Kristensen, B. (2013). Expression of the lysosomal-associated membrane protein-1 (LAMP-1) in astrocytomas. Int J Clin Exp Pathol, 6(7), 1294–1305.

Kadowaki, H., Nagai, A., Maruyama, T., Takami, Y., Satrimafitrah, P., Kato, H., Honda, A., Hatta, T., Natsume, T., Sato, T., Kai, H., Ichijo, H., & Nishitoh, H. (2015). Pre-emptive quality control protects the ER from protein overload via the proximity of ERAD components and SRP. Cell Reports, 13(5), 944–956. 10.1016/j.celrep.2015.09.047

Kim, S., Park, S. H., Kang, N., Ra, J. S., Myung, K., & Lee, K.-y. (2024). Polyubiquitinated PCNA triggers SLX4-mediated break-induced replication in alternat-ive lengthening of telomeres (ALT) cancer cells. Nucleic Acids Research, 52(19), 11785–11805. 10.1093/nar/gkae785

Kim, W. K., Kwon, Y., Jang, M., Park, M., Kim, J., Cho, S., Jang, D. G., Lee, W.-B., Jung, S. H., Choi, H. J., Min, B. S., Il Kim, T., Hong, S. P., Paik, Y.-K., & Kim, H. (2019). Β-catenin activation down-regulates cell-cell junction-related genes and induces epithelial-to-mesenchymal transition in colorectal cancers. Sci Rep, 9(1), 18440. 10.1038/s41598-019-54890-9

Kolberg, L., Raudvere, U., Kuzmin, I., Adler, P., Vilo, J., & Peterson, H. (2023). G:profiler—interoperable web service for functional enrichment analysis and gene identifier mapping (2023 update). Nucleic Acids Research, 51, W207–W212. 10.1093/nar/gkad347

Korotkevich, G., Sukhov, V., Budin, N., Shpak, B., Artyomov, M. N., & Sergushichev, A. (2016, June 20). Fast gene set enrichment analysis [Preprint, bioRxiv]. 10.1101/060012

Kravitz, S. N., Ferris, E., Love, M. I., Thomas, A., Quinlan, A. R., & Gregg, C. (2023). Random allelic expression in the adult human body. Cell Reports, 42(1), 111945. 10.1016/j.celrep.2022.111945

Kuhn, M. (2008). Building predictive models in *R* using the **caret** package. J. Stat. Soft., 28(5). 10.18637/jss.v028.i05

Kundra, R., Zhang, H., Sheridan, R., Sirintrapun, S. J., Wang, A., Ochoa, A., Wilson, M., Gross, B., Sun, Y., Madupuri, R., Satravada, B. A., Reales, D., Vakiani, E., Al-Ahmadie, H. A., Dogan, A., Arcila, M., Zehir, A., Maron, S., Berger, M. F., . . . Schultz, N. (2021). OncoTree: A cancer classification system for precision oncology. JCO Clinical Cancer Informatics, (5), 221–230. 10.1200/CCI.20.00108

Lee, S. G., Woo, S. M., Seo, S. U., Lee, C.-H., Baek, M.-C., Jang, S. H., Park, Z. Y., Yook, S., Nam, J.-O., & Kwon, T. K. (2024). Cathepsin d promotes polarization of tumor-associated macrophages and metastasis through TGFBI-CCL20 signaling. Exp Mol Med, 56(2), 383–394. 10.1038/s12276-024-01163-9

Li, Y., Dou, Y., Da Veiga Leprevost, F., Geffen, Y., Calinawan, A. P., Aguet, F., Akiyama, Y., Anand, S., Birger, C., Cao, S., Chaudhary, R., Chilappagari, P., Cieslik, M., Colaprico, A., Zhou, D. C., Day, C., Domagalski, M. J., Esai Selvan, M., Fenyö, D., . . . Chan, D. W. (2023). Proteogenomic data and resources for pan-cancer analysis. Cancer Cell, 41(8), 1397–1406. 10.1016/j.ccell.2023.06.009

Liberzon, A., Birger, C., Thorvaldsdóttir, H., Ghandi, M., Mesirov, J. P., & Tamayo, P. (2015). The molecular signatures database hallmark gene set collection. Cell Systems, 1(6), 417–425. 10.1016/j.cels.2015.12.004

Liu, Y., Borel, C., Li, L., Müller, T., Williams, E. G., Germain, P.-L., Buljan, M., Sajic, T., Boersema, P. J., Shao, W., Faini, M., Testa, G., Beyer, A., Antonarakis, S. E., & Aebersold, R. (2017). Systematic proteome and proteostasis profiling in human trisomy 21 fibroblast cells. Nat Commun, 8(1), 1212. 10.1038/s41467-017-01422-6

Lukow, D. A., & Sheltzer, J. M. (2022). Chromosomal instability and aneuploidy as causes of cancer drug resistance. Trends in Cancer, 8(1), 43–53. 10.1016/j.trecan.2021.09.002

Lundberg, S. M., & Lee, S.-I. (2017). A unified approach to interpreting model predictions [event-place: Long Beach, California, USA]. Proceedings of the 31st International Conference on Neural Information Processing Systems, 4768–4777. 10.48550/ARXIV.1705.07874

Mardin, B. R., Drainas, A. P., Waszak, S. M., Weischenfeldt, J., Isokane, M., Stütz, A. M., Raeder, B., Efthymiopoulos, T., Buccitelli, C., Segura-Wang, M., Northcott, P., Pfister, S. M., Lichter, P., Ellenberg, J., & Korbel, J. O. (2015). A cell-based model system links chromothripsis with hyperploidy. Molecular Systems Biology, 11(9), 828. 10.15252/msb.20156505

Mathieson, T., Franken, H., Kosinski, J., Kurzawa, N., Zinn, N., Sweetman, G., Poeckel, D., Ratnu, V. S., Schramm, M., Becher, I., Steidel, M., Noh, K.-M., Bergamini, G., Beck, M., Bantscheff, M., & Savitski, M. M. (2018). Systematic analysis of protein turnover in primary cells. Nat Commun, 9(1), 689. 10.1038/s41467-018-03106-1

McShane, E., Sin, C., Zauber, H., Wells, J. N., Donnelly, N., Wang, X., Hou, J., Chen, W., Storchova, Z., Marsh, J. A., Valleriani, A., & Selbach, M. (2016). Kinetic analysis of protein stability reveals age-dependent degradation. Cell, 167(3), 803–815.e21. 10.1016/j.cell.2016.09.015

Mertins, P., Mani, D. R., Ruggles, K. V., Gillette, M. A., Clauser, K. R., Wang, P., Wang, X., Qiao, J. W., Cao, S., Petralia, F., Kawaler, E., Mundt, F., Krug, K., Tu, Z., Lei, J. T., Gatza, M. L., Wilkerson, M., Perou, C. M., Yellapantula, V., . . . NCI CPTAC. (2016). Proteogenomics connects somatic mutations to signalling in breast cancer. Nature, 534(7605), 55–62. 10.1038/nature18003

Meyers, R. M., Bryan, J. G., McFarland, J. M., Weir, B. A., Sizemore, A. E., Xu, H., Dharia, N. V., Montgomery, P. G., Cowley, G. S., Pantel, S., Goodale, A., Lee, Y., Ali, L. D., Jiang, G., Lubonja, R., Harrington, W. F., Strickland, M., Wu, T., Hawes, D. C., . . . Tsherniak, A. (2017). Computational correction of copy number effect improves specificity of CRISPR–cas9 essentiality screens in cancer cells. Nat Genet, 49(12), 1779–1784. 10.1038/ng.3984

Moldovan, G.-L., Pfander, B., & Jentsch, S. (2007). PCNA, the maestro of the replication fork. Cell, 129(4), 665–679. 10.1016/j.cell.2007.05.003

Motegi, A., Liaw, H.-J., Lee, K.-Y., Roest, H. P., Maas, A., Wu, X., Moinova, H., Markowitz, S. D., Ding, H., Hoeijmakers, J. H. J., & Myung, K. (2008). Polyubiquitination of proliferating cell nuclear antigen by HLTF and SHPRH prevents genomic in-stability from stalled replication forks. Proc. Natl. Acad. Sci. U.S.A., 105(34), 12411–12416. 10.1073/pnas.0805685105

Mounir, M., Lucchetta, M., Silva, T. C., Olsen, C., Bontempi, G., Chen, X., Noushmehr, H., Colaprico, A., & Papaleo, E. (2019). New functionalities in the TCGAbiolinks package for the study and integration of cancer data from GDC and GTEx (E. Wang, Ed.). PLoS Comput Biol, 15(3), e1006701. 10.1371/journal.pcbi.1006701

Mueller, S. C., Ghersi, G., Akiyama, S. K., Sang, Q.-X. A., Howard, L., Pineiro-Sanchez, M., Nakahara, H., Yeh, Y., & Chen, W.-T. (1999). A novel protease-docking function of integrin at invadopodia. Journal of Biological Chemistry, 274(35), 24947– 24952. 10.1074/jbc.274.35.24947

Muenzner, J., Trébulle, P., Agostini, F., Zauber, H., Messner, C. B., Steger, M., Kilian, C., Lau, K., Barthel, N., Lehmann, A., Textoris-Taube, K., Caudal, E., Egger, A.-S., Amari, F., De Chiara, M., Demichev, V., Gossmann, T. I., Mülleder, M., Liti, G., . . . Ralser, M. (2024). Natural proteome diversity links aneuploidy tolerance to protein turnover. Nature, 630(8015), 149–157. 10.1038/s41586-024-07442-9

Nusinow, D. P., Szpyt, J., Ghandi, M., Rose, C. M., McDonald, E. R., Kalocsay, M., Jané-Valbuena, J., Gelfand, E., Schweppe, D. K., Jedrychowski, M., Golji, J., Porter, D. A., Rejtar, T., Wang, Y. K., Kryukov, G. V., Stegmeier, F., Erickson, B. K., Garraway, L. A., Sellers, W. R., & Gygi, S. P. (2020). Quantitative proteomics of the cancer cell line encyclopedia. Cell, 180(2), 387–402.e16. 10.1016/j.cell.2019.12.023

Oh, S., Abdelnabi, J., Al-Dulaimi, R., Aggarwal, A., Ramos, M., Davis, S., Riester, M., & Waldron, L. (2022). HGNChelper: Identification and correction of invalid gene symbols for human and mouse. F1000Res, 9, 1493. 10.12688/f1000research.28033.2

Oromendia, A. B., & Amon, A. (2014). Aneuploidy: Implications for protein homeostasis and disease. Disease Models & Mechanisms, 7(1), 15–20. 10.1242/dmm.013391

Oromendia, A. B., Dodgson, S. E., & Amon, A. (2012). Aneuploidy causes proteotoxic stress in yeast. Genes Dev., 26(24), 2696–2708. 10.1101/gad.207407.112

Pacini, C., Dempster, J. M., Boyle, I., Gonçalves, E., Najgebauer, H., Karakoc, E., Van Der Meer, D., Barthorpe, A., Lightfoot, H., Jaaks, P., McFarland, J. M., Garnett, M. J., Tsherniak, A., & Iorio, F. (2021). Integrated cross-study datasets of genetic dependencies in cancer. Nat Commun, 12(1), 1661. 10.1038/s41467-021-21898-7

Passerini, V., Ozeri-Galai, E., De Pagter, M. S., Donnelly, N., Schmalbrock, S., Kloosterman, W. P., Kerem, B., & Storchová, Z. (2016). The presence of extra chromosomes leads to genomic instability. Nat Commun, 7(1), 10754. 10.1038/ncomms10754

Piovesan, D., Necci, M., Escobedo, N., Monzon, A. M., Hatos, A., Mičetić, I., Quaglia, F., Paladin, L., Ramasamy, P., Dosztányi, Z., Vranken, W. F., Davey, N. E., Parisi, G., Fuxreiter, M., & Tosatto, S. C. E. (2021). MobiDB: Intrinsically disordered proteins in 2021. Nucleic Acids Research, 49, D361–D367. 10.1093/nar/gkaa1058

Raine, K. M., Van Loo, P., Wedge, D. C., Jones, D., Menzies, A., Butler, A. P., Teague, J. W., Tarpey, P., Nik-Zainal, S., & Campbell, P. J. (2016). ascatNgs: Identifying somatically acquired copy-number alterations from whole-genome sequencing data. CP in Bioinformatics, 56(1). 10.1002/cpbi.17

Redelmeier, A., Jullum, M., & Aas, K. (2020, July 2). Explaining predictive models with mixed features using shapley values and conditional inference trees [Preprint]. Retrieved January 6, 2024, from http://arxiv.org/abs/2007.01027

Replogle, J. M., Zhou, W., Amaro, A. E., McFarland, J. M., Villalobos-Ortiz, M., Ryan, J., Letai, A., Yilmaz, O., Sheltzer, J., Lippard, S. J., Ben-David, U., & Amon, A. (2020). Aneuploidy increases resistance to chemotherapeutics by antagonizing cell division. Proc. Natl. Acad. Sci. U.S.A., 117(48), 30566–30576. 10.1073/pnas.2009506117

Ritchie, M. E., Phipson, B., Wu, D., Hu, Y., Law, C. W., Shi, W., & Smyth, G. K. (2015). Limma powers differential expression analyses for RNA-sequencing and microarray studies. Nucleic Acids Research, 43(7), e47–e47. 10.1093/nar/gkv007

Robin, X., Turck, N., Hainard, A., Tiberti, N., Lisacek, F., Sanchez, J.-C., & Müller, M. (2011). pROC: An open-source package for r and s+ to analyze and compare ROC curves. BMC Bioinformatics, 12(1), 77. 10.1186/1471-2105-12-77

Rochefort, H., Capony, F., & Garcia, M. (1990). Cathepsin d: A protease involved in breast cancer metastasis. Cancer Metast Rev, 9(4), 321–331. 10.1007/BF00049522

Ruan, J., Zheng, H., Rong, X., Rong, X., Zhang, J., Fang, W., Zhao, P., & Luo, R. (2016). Over-expression of cathepsin b in hepatocellular carcinomas predicts poor prognosis of HCC patients. Mol Cancer, 15, 17. 10.1186/s12943-016-0503-9

Santaguida, S., & Amon, A. (2015). Short- and long-term effects of chromosome mis-segregation and aneuploidy. Nat Rev Mol Cell Biol, 16(8), 473–485. 10.1038/nrm4025

Sarafian, V., Jadot, M., Foidart, J.-M., Letesson, J.-J., Van Den Brûle, F., Castronovo, V., Wattiaux, R., & Wattiaux-De Coninck, S. (1998). Expression of lamp-1 and lamp-2 and their interactions with galectin-3 in human tumor cells. Int. J. Cancer, 75(1), 105–111. 10.1002/(SICI)1097-0215(19980105)75:1<105::AID-IJC16>3.0.CO;2-F

Schukken, K. M., & Sheltzer, J. M. (2022). Extensive protein dosage compensation in aneuploid human cancers. Genome Res., 32(7), 1254–1270. 10.1101/gr.276378.121

Sdeor, E., Okada, H., Saad, R., Ben-Yishay, T., & Ben-David, U. (2024). Aneuploidy as a driver of human cancer. Nat Genet, 56(10), 2014–2026. 10.1038/s41588-024-01916-2

Senger, G., & Schaefer, M. H. (2021). Protein complex organization imposes constraints on proteome dysregulation in cancer. Front. Bioinform., 1, 723482. 10.3389/fbinf.2021.723482

Sheltzer, J. M., Torres, E. M., Dunham, M. J., & Amon, A. (2012). Transcriptional consequences of aneuploidy. Proc. Natl. Acad. Sci. U.S.A., 109(31), 12644–12649. 10.1073/pnas.1209227109

Shih, J., Sarmashghi, S., Zhakula-Kostadinova, N., Zhang, S., Georgis, Y., Hoyt, S. H., Cuoco, M. S., Gao, G. F., Spurr, L. F., Berger, A. C., Ha, G., Rendo, V., Shen, H., Meyerson, M., Cherniack, A. D., Taylor, A. M., & Beroukhim, R. (2023). Cancer aneuploidies are shaped primarily by effects on tumour fitness. Nature, 619(7971), 793–800. 10.1038/s41586-023-06266-3

Shima, N., Alcaraz, A., Liachko, I., Buske, T. R., Andrews, C. A., Munroe, R. J., Hartford, S. A., Tye, B. K., & Schimenti, J. C. (2007). A viable allele of mcm4 causes chromosome instability and mammary adenocarcinomas in mice. Nat Genet, 39(1), 93–98. 10.1038/ng1936

Shukla, A., Nguyen, T. H. M., Moka, S. B., Ellis, J. J., Grady, J. P., Oey, H., Cristino, A. S., Khanna, K. K., Kroese, D. P., Krause, L., Dray, E., Fink, J. L., & Duijf, P. H. G. (2020). Chromosome arm aneuploidies shape tumour evolution and drug response. Nat Commun, 11(1), 449. 10.1038/s41467-020-14286-0

Silva, T. C., Colaprico, A., Olsen, C., D’Angelo, F., Bontempi, G., Ceccarelli, M., & Noushmehr, H. (2016). TCGA workflow: Analyze cancer genomics and epigenomics data using bioconductor packages. F1000Res, 5, 1542. 10.12688/f1000research.8923.2

Soto, M., Raaijmakers, J. A., Bakker, B., Spierings, D. C., Lansdorp, P. M., Foijer, F., & Medema, R. H. (2017). P53 prohibits propagation of chromosome segregation errors that produce structural aneuploidies. Cell Reports, 19(12), 2423–2431. 10.1016/j.celrep.2017.05.055

Sousa, A., Gonçalves, E., Mirauta, B., Ochoa, D., Stegle, O., & Beltrao, P. (2019). Multi-omics characterization of interaction-mediated control of human protein abundance levels. Molecular & Cellular Proteomics, 18(8), S114–S125. 10.1074/mcp.RA118.001280

Stingele, S., Stoehr, G., Peplowska, K., Cox, J., Mann, M., & Storchova, Z. (2012). Global analysis of genome, transcriptome and proteome reveals the response to aneuploidy in human cells. Mol Syst Biol, 8(1), 608. 10.1038/msb.2012.40

Subramanian, A., Tamayo, P., Mootha, V. K., Mukherjee, S., Ebert, B. L., Gillette, M. A., Paulovich, A., Pomeroy, S. L., Golub, T. R., Lander, E. S., & Mesirov, J. P. (2005). Gene set enrichment analysis: A knowledge-based approach for interpreting genome-wide expression profiles. Proc. Natl. Acad. Sci. U.S.A., 102(43), 15545– 15550. 10.1073/pnas.0506580102

Suehnholz, S. P., Nissan, M. H., Zhang, H., Kundra, R., Nandakumar, S., Lu, C., Carrero, S., Dhaneshwar, A., Fernandez, N., Xu, B. W., Arcila, M. E., Zehir, A., Syed, A., Brannon, A. R., Rudolph, J. E., Paraiso, E., Sabbatini, P. J., Levine, R. L., Dogan, A., . . . Chakravarty, D. (2024). Quantifying the expanding landscape of clinical actionability for patients with cancer. Cancer Discovery, 14(1), 49–65. 10.1158/2159-8290.CD-23-0467

Swanton, E., High, S., & Woodman, P. (2003). Role of calnexin in the glycan-independent quality control of proteolipid protein. EMBO J, 22(12), 2948–2958. 10.1093/emboj/cdg300

Szklarczyk, D., Kirsch, R., Koutrouli, M., Nastou, K., Mehryary, F., Hachilif, R., Gable, A. L., Fang, T., Doncheva, N. T., Pyysalo, S., Bork, P., Jensen, L. J., & von Mering, C. (2023). The STRING database in 2023: Protein–protein association networks and functional enrichment analyses for any sequenced genome of interest. Nucleic Acids Research, 51, D638–D646. 10.1093/nar/gkac1000

Taggart, J. C., Zauber, H., Selbach, M., Li, G.-W., & McShane, E. (2020). Keeping the proportions of protein complex components in check. Cell Systems, 10(2), 125–132. 10.1016/j.cels.2020.01.004

Tang, Y.-C., Williams, B. R., Siegel, J. J., & Amon, A. (2011). Identification of aneuploidy-selective antiproliferation compounds. Cell, 144(4), 499–512. 10.1016/j.cell.2011.01.017

Taylor, A. M., Shih, J., Ha, G., Gao, G. F., Zhang, X., Berger, A. C., Schumacher, S. E., Wang, C., Hu, H., Liu, J., Lazar, A. J., Cherniack, A. D., Beroukhim, R., Meyerson, M., Caesar-Johnson, S. J., Demchok, J. A., Felau, I., Kasapi, M., Ferguson, M. L., . . . Mariamidze, A. (2018). Genomic and functional approaches to understanding cancer aneuploidy. Cancer Cell, 33(4), 676–689.e3. 10.1016/j.ccell.2018.03.007

Torres, E. M., Dephoure, N., Panneerselvam, A., Tucker, C. M., Whittaker, C. A., Gygi, S. P., Dunham, M. J., & Amon, A. (2010). Identification of aneuploidy-tolerating mutations. Cell, 143(1), 71–83. 10.1016/j.cell.2010.08.038

Torres, E. M., Sokolsky, T., Tucker, C. M., Chan, L. Y., Boselli, M., Dunham, M. J., & Amon, A. (2007). Effects of aneuploidy on cellular physiology and cell division in haploid yeast. Science, 317(5840), 916–924. 10.1126/science.1142210

Turakhiya, A., Meyer, S. R., Marincola, G., Böhm, S., Vanselow, J. T., Schlosser, A., Hofmann, K., & Buchberger, A. (2018). ZFAND1 recruits p97 and the 26s proteasome to promote the clearance of arsenite-induced stress granules. Molecular Cell, 70(5), 906–919.e7. 10.1016/j.molcel.2018.04.021

UCSC Genome Bioinformatics Group. (n.d.). UCSC human genome browser [DATASET]. Retrieved December 24, 2023, from https://genome.ucsc.edu/cgi-bin/hgTables

Unk, I., Hajdú, I., Fátyol, K., Hurwitz, J., Yoon, J.-H., Prakash, L., Prakash, S., & Haracska, L. (2008). Human HLTF functions as a ubiquitin ligase for proliferating cell nuclear antigen polyubiquitination. Proc. Natl. Acad. Sci. U.S.A., 105(10), 3768–3773. 10.1073/pnas.0800563105

Upadhya, S. R., & Ryan, C. J. (2022). Experimental reproducibility limits the correlation between mRNA and protein abundances in tumor proteomic profiles. Cell Reports Methods, 2(9), 100288. 10.1016/j.crmeth.2022.100288

Viganó, C., von Schubert, C., Ahrné, E., Schmidt, A., Lorber, T., Bubendorf, L., De Vetter, J. R. F., Zaman, G. J. R., Storchova, Z., & Nigg, E. A. (2018). Quantitative proteomic and phosphoproteomic comparison of human colon cancer DLD-1 cells differing in ploidy and chromosome stability (F. A. Barr, Ed.). MBoC, 29(9), 1031– 1047. 10.1091/mbc.E17-10-0577

Yang, E., Van Nimwegen, E., Zavolan, M., Rajewsky, N., Schroeder, M., Magnasco, M., & Darnell, J. E. (2003). Decay rates of human mRNAs: Correlation with functional characteristics and sequence attributes. Genome Res., 13(8), 1863–1872. 10.1101/gr.1272403

Zhang, B., Wang, J., Wang, X., Zhu, J., Liu, Q., Shi, Z., Chambers, M. C., Zimmerman, L. J., Shaddox, K. F., Kim, S., Davies, S. R., Wang, S., Wang, P., Kinsinger, C. R., Rivers, R. C., Rodriguez, H., Townsend, R. R., Ellis, M. J. C., Carr, S. A., . . . the NCI CPTAC. (2014). Proteogenomic characterization of human colon and rectal cancer. Nature, 513(7518), 382–387. 10.1038/nature13438

Zhang, H., Liu, T., Zhang, Z., Payne, S. H., Zhang, B., McDermott, J. E., Zhou, J.-Y., Petyuk, V. A., Chen, L., Ray, D., Sun, S., Yang, F., Chen, L., Wang, J., Shah, P., Cha, S. W., Aiyetan, P., Woo, S., Tian, Y., . . . Townsend, R. R. (2016). Integrated proteogenomic characterization of human high-grade serous ovarian cancer. Cell, 166(3), 755–765. 10.1016/j.cell.2016.05.069

